# Got Milk? Maternal immune activation during the mid-lactational period affects nutritional milk quality and adolescent offspring sensory processing in male and female rats

**DOI:** 10.1101/2022.01.03.474857

**Authors:** Holly DeRosa, Salvatore G. Caradonna, Hieu Tran, Jordan Marrocco, Amanda C. Kentner

## Abstract

Previous studies have underscored the importance of breastfeeding and parental care on offspring development and behavior. However, their contribution as dynamic variables in animal models of early life stress are often overlooked. In the present study, we investigated how lipopolysaccharide (LPS)-induced maternal immune activation (MIA) on postnatal day (P)10 affects maternal care, milk, and offspring development. MIA was associated with elevated milk corticosterone concentrations on P10, which recovered by P11. In contrast, both milk triglyceride and percent creamatocrit values demonstrated a prolonged decrease following inflammatory challenge. Adolescent MIA offspring were heavier, which is often suggestive of poor early life nutrition. While MIA did not decrease maternal care quality, there was a significant compensatory increase in maternal licking and grooming the day following inflammatory challenge. However, this did not protect against disrupted neonatal huddling or later-life alterations in sensorimotor gating, conditioned fear, mechanical allodynia, or reductions in hippocampal parvalbumin expression in MIA offspring. MIA-associated changes in brain and behavior were likely driven by differences in milk nutritional values and not by direct exposure to LPS or inflammatory molecules as neither LPS binding protein nor interleukin-6 milk levels differed between groups. These findings reflected comparable microbiome and transcriptomic patterns at the genome-wide level. Animal models of early life stress can impact both parents and their offspring. One mechanism that can mediate the effects of such stressors is changes to maternal lactation quality which our data show can confer multifaceted and compounding effects on offspring physiology and behavior.

## Introduction

Epidemiological studies have shown that prenatal exposure to maternal immune activation (MIA) is a leading risk factor for developing psychiatric disorders such as autism and schizophrenia (1-3). Administering immunogens such as lipopolysaccharide (LPS) or polyinosinic:polycytidylic acid (poly I:C) to pregnant animals induces MIA by promoting a cascade of inflammatory cytokines and activating the hypothalamic-pituitary adrenal (HPA) axis of the mother (4-7). Together, these physiological responses are recognized to affect the long-term development of offspring (4, 7). However, the impact on mothers and their offspring when MIA occurs during lactation is relatively unexplored. This is surprising because infections during the postnatal period are common, particularly in low resource settings, and can result in severe morbidity and mortality (8-11).

LPS administered to rodent dams during the lactational period can impact maternal care by reducing the amount of time spent in efficient arched back nursing postures and the number of pups retrieved and returned back to the nest when displaced (12). In line with the behavioral alterations, MIA in lactating dams may influence maternal milk quality. For example, maternal LPS treatment on postnatal days 4, 11, and 18 reduced the mRNA expression of milk nutrient precursors such as glucose and fatty acid transporters in the rat mammary gland (13). Exposure to MIA during the lactational period can also impact offspring behavior. For example, Nascimento and colleagues (14) observed a reduction in ultrasonic vocalizations in male neonates shortly after their dams were treated with LPS. Although this presumably involved changes in various forms of maternal input signals (e.g., parental care and/or milk quality), the mechanisms of this MIA-derived effect on offspring behavior remain unclear. While there is a small amount of evidence suggesting how breastfeeding can transmit certain pathogenic bacteria and viral infections that lead to infant disease (15-17), to our knowledge no epidemiological study has assessed how maternal infection may impact the trajectory of infant development.

Other exogenous factors have been shown to alter contents in breastmilk and subsequent offspring development in animals and humans (18-22). For example, psychological stress in humans reduces milk microbial diversity (23). Further, cortisol in maternal milk was found to predict temperament and promote greater body weights in infant primates, although milk energy content was also associated with body weight (24). Together, this evidence suggests that 1) disturbances to the maternal-neonatal environment alters the developmental trajectory of offspring, and 2) these alterations are mediated by changes in maternal care and/or nutritional milk quality. To explore these ideas further, we assessed the effects of MIA challenge during the mid-lactational period on maternal care, maternal milk composition, and offspring brain development and behavior in Sprague-Dawley rats. While a majority of studies have focused on adult behavioral outcomes following gestational MIA exposure, longitudinal evidence suggests that differences in MIA-associated alterations in brain development emerge in adolescence (25-26). Therefore, we curated a battery of behavioral measures with proven translational potential (27-28) to uncover how the developing brain may be disrupted by maternal infection. We also examined hippocampal expression of parvalbumin given the role of this critical neuronal population in the pathophysiology of neuropsychiatric disorders in humans and in animal models of MIA (29).

## Methods and Experimental Overview

All animal procedures were performed in accordance with the Association for Assessment and Accreditation of Laboratory Animal Care with protocols approved by the Massachusetts College of Pharmacy and Health Sciences Institutional Animal Care and Use Committee. **Figure 1A** and Table 1 outline the experimental procedures and groups evaluated in this study. Please see the **Supplementary Methods** and **Supplementary Table 1** for a more detailed description of the methodological protocols and statistical analyses used in this study. Briefly, on postnatal day 10 (P10; equivalent to the mature milk phase), Sprague-Dawley rat mothers were removed from their litters and placed into a clean cage located in a separate procedure room. Dams were challenged intraperitoneally (i.p.) with either 100 μg/kg of the inflammatory endotoxin, lipopolysaccharide (LPS; Escherichia coli, serotype 026:B6; L-3755), or pyrogen-free saline between 08:00hrs and 10:00hrs. During this period the offspring stayed in their regular holding room and were placed into a smaller clean cage positioned on top of a heating pad to facilitate body temperature maintenance. Offspring body weights were collected immediately before returning to their dams and again 2 and 24 hours later, alongside the inspection of milk bands, to monitor health. Exactly 2 hours after receiving either a saline or LPS injection, dams were anesthetized with isoflurane in O2 and administered 0.2 mL of oxytocin (20 USP/mL i.p.) to facilitate milk production. The pharmacokinetic profile of isoflurane suggests it is not absorbed by offspring and that breastfeeding can resume immediately after anesthesia (30-31). Similarly, oxytocin is not expected to affect offspring given its short plasma half-life of 1-6 minutes which is reduced further during lactation (32). Teats were prepared by moistening the collection areas with distilled water. Milk was collected by gently squeezing the base of each teat, and 20uL of expelled milk was immediately processed to measure levels of creamatocrit while an additional ∼500uL was stored at -80°C for additional processing. On P11, dams were separated from their litters again for 2 hours prior to milk collection which occurred 24 hours after LPS or saline treatment (see 33).

**Table 1.**
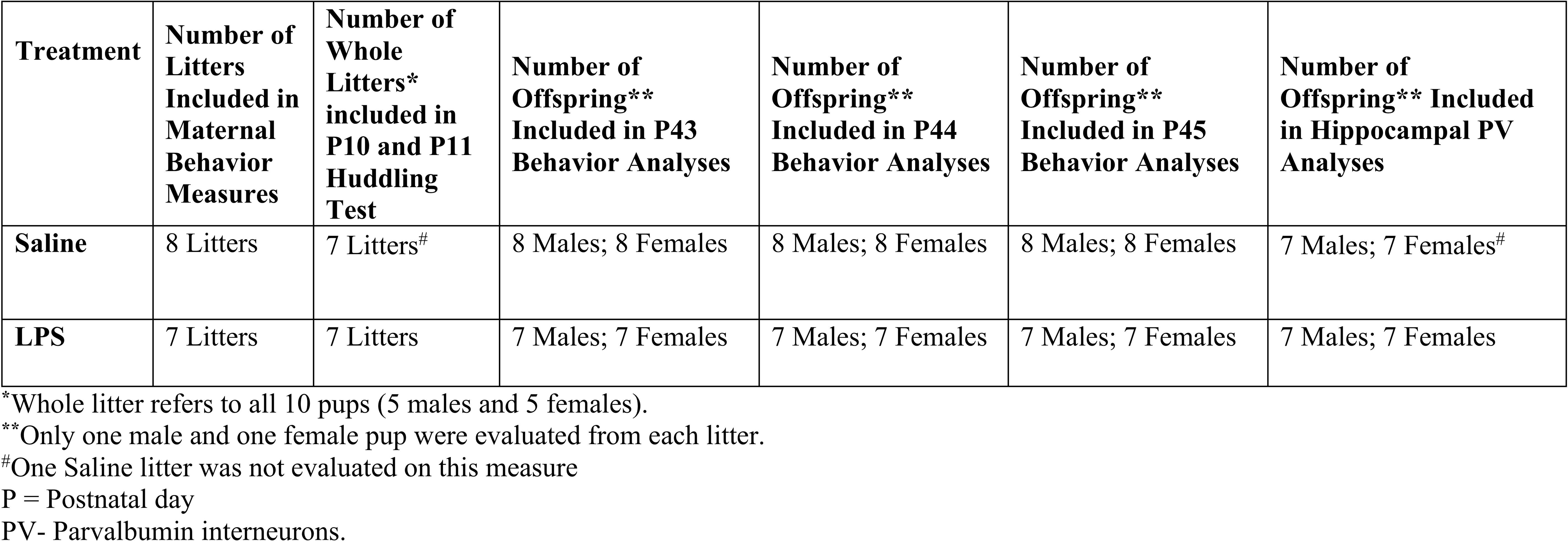
Litter Treatments and Endpoints.

**Figure 1.**
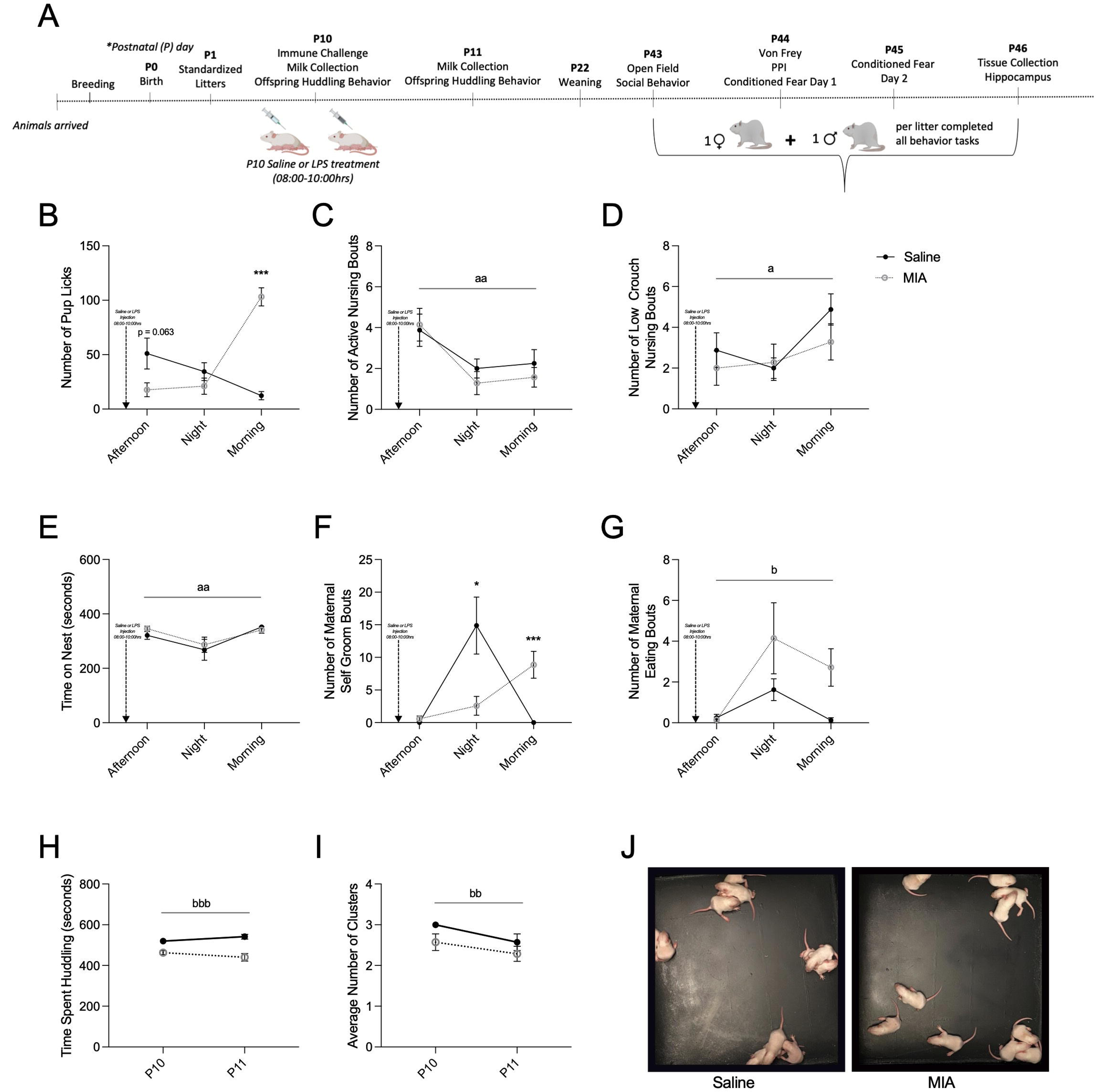
Maternal care and offspring huddling behaviors following maternal immune activation (MIA) during the mid-lactational period. (A) Timeline of experimental procedures. Maternal care was evaluated in the afternoon and night of postnatal day (P) 10 and the morning of P11. (B) MIA x Time interaction for the total number of pup licks (F(2, 26) = 31.435, p = 0.001, *n*_*p*_^2^ = 0.701). Pup licking and grooming were elevated in MIA dams on P11 (p = 0.001). The circadian cycle influenced the display of (C) active nursing bouts (F(2, 26) = 7.752, p = 0.0022, *n*_*p*_^2^ = 0.374), which were elevated in both groups on P10. (D) Low crouch nursing bouts (F(2, 26) = 3.960, p = 0.032, *n*_*p*_^2^ = 0.233), and (E) total time (seconds) that the dam spent on nest were also affected by the circadian cycle, as dams spent less time in the low crouch nursing posture and overall less time on the nest during the evening on P10. Total number of (F) self-groom bouts demonstrated a MIA x Time interaction (F(1.322, 26) = 12.33, p = 0.001, *n*_*p*_^2^ = 0.485); Huyndt Feldt correction. MIA was associated with a reduction in self-grooming bouts on the evening of P10 (p = 0.025), which were increased on the morning of P11 (p = 0.0001). A main effect of MIA was observed for (G) the number of eating bouts (F(1, 13) = 10.64, p = 0.006, *n*_*p*_^2^ = 0.450). (H) Total time pups spent huddling (seconds) was reduced by MIA (F(1, 12) = 30.079, p = 0.001, *n*_*p*_^2^ = 0.715) as was the (I) average number of pup clusters across P10 and P11 (F(1, 12) = 8.33, p = 0.014, *n*_*p*_^2^ = 0.410). (J) Shows representative photographs of the huddling behavior of saline and MIA litters. Data are expressed as mean ± SEM; Saline: n = 8; MIA: n = 7. *p < 0.05, ***p <0.001, MIA versus Saline; ^a^p < 0.05, ^aa^p < 0.01, main effect of time; ^b^p < 0.05, ^bb^p < 0.01, ^bbb^p < 0.001, main effect of MIA.

Milk quality was determined through the use of several assays (see 20, 33) to measure levels of milk creamatocrit, lactose (Sigma-Aldrich Cat. #MAK017), triglycerides (Abcam, Cat. #ab65336), protein (Thermo Fisher Scientific, Cat. #23227), corticosterone (#ADI-900– 097, Enzo Life Sciences, Farmingdale, NY), immunoglobulin (Ig) A (Bethyl Laboratories, Cat. #E111-102), interleukin (IL)-6 (Thermo Fisher Scientific, Cat. #BMS625) and lipopolysaccharide binding protein (LBP; Abcam, Cat. #ab269542). Commercially available immunoassays were performed according to the manufacturer’s instructions.

The milk microbiome was examined by sequencing the V3-V4 regions of the 16S gene (Zymo Research, Irvine, CA). Following preparation, the final library was quantified using TapeStation® (Agilent Technologies, Santa Clara, CA) and Qubit® (Thermo Fisher Scientific, Waltham, WA), and sequenced using an Illumina® MiSeq™ with a v3 reagent kit (600 cycles) and 10% PhiX spike-in. Taxonomic profiling of was performed with Uclust (Qiime v.1.9.1) and the Zymo Research Database (Zymo Research, Irvine, CA).

Following RNA-isolation (see 20, 33; QIAGEN, Cat. #217004), milk samples were more broadly analyzed using RNA Sequencing (RNA-seq). After determining fragment size and concentration using TapeStation® (Agilent Technologies, Santa Clara, CA), an Illumina NovaSeq 6000 was used to obtain 100-bp reads and samples were read at a sequencing depth of approximately 50 million reads.

Maternal behavior was assessed in the afternoon (15:00-17:00 hrs) and evening (20:00-22:00 hrs) of P10 and in the morning (07:30-9:30 hrs) on P11. Dams were evaluated for 1-minute intervals over 6 observation periods on behaviors including the frequency of pup-directed behaviors, self-directed behaviors, nest building/digging behavior, and the total time spent on the nest. The offspring of saline and MIA treated dams completed an assessment of neonatal huddling, a form of social thermoregulation, 2 hours after being reunited with their mother on P10 and P11. Starting on P43, one male and one female offspring from each litter were assessed in several behavioral measures including the open field and social preference tasks, as well as assessments of mechanical allodynia (the von Frey test), sensorimotor gating (prepulse inhibition; PPI), and conditioned fear (see 34-37). On P46, whole hippocampus was collected and stored at -80°C for future processing.

To measure hippocampal parvalbumin (PV) expression using western blotting, 20μg of protein was loaded into each well of MiniProtean® gels (Bio Rad Laboratories, Cat. #4568101). Gels were transferred onto nitrocellulose membranes (Bio Rad Laboratories, Cat. #1620147) and blocked in 5% nonfat milk with TBS + 0.05% Tween 20 (TBST) for 1 hour at room temperature. Membranes were then washed with TBST and incubated in a 1:1000 dilution of parvalbumin (PV) antibody (RnD Systems, Cat. # AF5058) plus TBS overnight at 4°C. The next morning, membranes were washed and incubated in an HRP-conjugated secondary antibody (1:1000, RnD Systems, Cat., #HAF016) made in 1% nonfat milk with TBS for 1 hour at room temperature. Membranes were then washed and exposed to a chemiluminescent substrate (Thermo Fisher Scientific, Cat. #34580) for 5 minutes prior to being scanned. After imaging, membranes were stripped (Thermo Fisher, Cat. # 21062) for 15 minutes at 37°C, blocked, washed, and incubated in beta actin primary antibody (1:1000, Thermo Fisher Scientific, Cat. #MA515739) for 1 hour at room temperature. Membranes were exposed again to the chemiluminescence substrate and imaged. Densitometry measures were used to obtain a ratio of PV/β-actin in order to quantify differences between groups.

Two-way repeated measure ANOVAs (MIA x Time) were used to evaluate milk and behavioral measures across P10 and P11. Violations to the assumption of sphericity were addressed using the Huyndt Feldt correction. One-way ANOVAs were used as appropriate for all other measures unless there were violations to the assumption of normality (Shapiro-Wilk test) in which case Kruskal-Wallis tests were employed (expressed as Χ^*2*^). The mechanical allodynia threshold data was assessed using body weight as a covariate. The partial eta-squared (*n*_*p*_^2^) is reported as an index of effect size for the ANOVAs (38). Because the dataset was not powered to evaluate sex-differences directly, male and female animals were evaluated separately (39). All data are expressed as mean ± SEM. Offspring data are depicted as each sex separately and collapsed across males and females for display purposes.

For microbiome sequencing, samples underwent composition visualization, in addition to alpha-diversity and beta-diversity analyses (Qiime v.1.9.1) and statistical comparisons were performed using Kruskal-Wallis (40). Linear discriminant analysis effect size (LEfSe; http://huttenhower.sph.harvard.edu/lefse/) was utilized to determine significant differences in taxonomy abundance between each group as previously described (41-42).

For RNA-Seq, DESeq2 was used to determine differentially expressed genes based on a p < 0.05, Benjamini–Hochberg false discovery rate corrected (FDR) and fold change (FC) > 1.3 (43). Heatmaps were generated using the MultiExperiment Viewer (National Library of Medicine, USA), while gene ontology was determined using the Database for Annotation, Visualization and Integrated Discovery functional annotation cluster tool (https://david.ncifcrf.gov/).

## Results

### MIA challenge affected maternal behavior and offspring huddling

Immune challenge during lactation significantly impacted the number of maternal care bouts directed towards pups. For example, a MIA by time interaction was identified for the number of licks pups received following maternal LPS exposure (**Figure 1B**). While the number of licks received was not significantly affected on P10, there was a general pattern of reduced care provided by MIA treated dams on the afternoon of P10. This was followed by a rebound of maternal pup licking by MIA dams on the morning of P11, possibly to compensate for overall reduced maternal care due to illness. The display of active and low crouch nursing behaviors and time on nest were affected by the circadian cycle (**Figure 1C, D, E**), but only time on nest was impacted by MIA experience. Maternal self-directed care was impacted as a function of sickness. There was a significant MIA by time interaction for total number of self-grooms (**Figure 1F**); MIA dams had fewer bouts of grooming on the evening of P10, which reversed substantially by the morning of P11. Similarly, the number of maternal eating bouts was elevated on P11, although there were no significant effects of time (**Figure 1G**) there was a main effect of MIA (**Figure 1G**). Additional maternal care data are presented in **Supplementary Figure 1A-D**.

Huddling behavior was significantly reduced in the offspring of MIA dams compared to saline (main effect of MIA; time spent huddling, **Figure 1H**; average number of clusters: **Figure 1I)**. See **Figure 1J** for representative displays of huddling behavior in saline and MIA offspring.

### MIA challenge impacted nutritional composition of milk and microbiome communities

For milk collection (see photographs in **Figure 2A** depicting the milk collection procedure), there was a significant MIA by time interaction for maternal milk levels of corticosterone. This stress-associated hormone was elevated in milk samples 2-hours following MIA challenge and was recovered by P11 (**Figure 2B)**. A MIA by time interaction was also present for % creamatocrit (**Figure 2C**), which is linearly related to the fat concentration and energy content of milk (44, 45; see **Supplementary Figure 2A, B**). These measures were significantly lower in the milk of MIA exposed mothers compared to saline on P11. Triglycerides were also reduced in the milk of MIA treated dams on P11 (**Figure 2D**).

**Figure 2.**
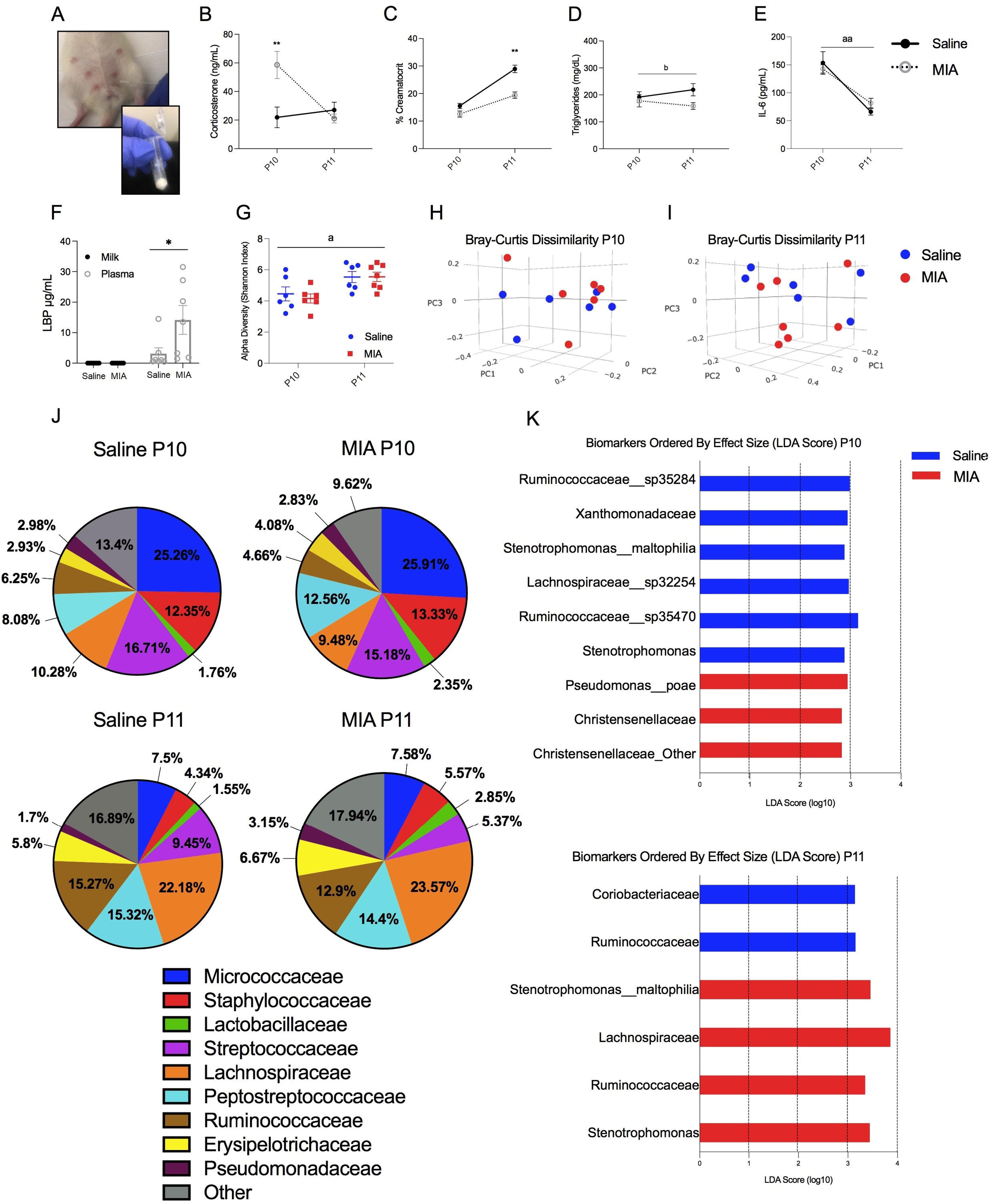
Nutritional profile and microbiome community distribution of milk following maternal immune activation (MIA) during the mid-lactational period. (A) Photographs of maternal milk collection. A significant MIA x Time interaction was observed for maternal milk levels (Saline: n = 8; MIA: n = 7) of (B) corticosterone (ng/mL; F(1, 13) = 8.496, p = 0.013, *n*_*p*_^2^ = 0.415), which was elevated in the milk of MIA dams on P10 (p = 0.009). A MIA x Time interaction was observed for (C) percent (%) creamatocrit (F(1, 13) = 5.86, p = 0.032, *n*_*p*_^2^ = 0.858), which was significantly lower in MIA samples on P11 (p = 0.002). A significant main effect of MIA was observed for (D) milk triglycerides (mg/dL; F(1, 13) = 6.496, p = 0.026, *n*_*p*_^2^ = 0.351). There was a main effect of time on (E) milk interleukin (IL)-6 (pg/mL) concentrations (F(1,13) = 33.01, p < 0.001, *n*_*p*_^2^ = 0.717) in that IL-6 was increased in both groups on P11. Levels of (F) LPS binding protein (LPB; ug/mL) were increased in the blood plasma of MIA dams (*X*^2^(1)=5.00, p=0.025), but there were negligible levels in maternal milk (p>0.05). (G) Alpha diversity of P10 and P11 milk samples along the Shannon Index was increased in both groups on P11 compared to P10 (*X*^2^(1) = 5.33, p = 0.021). (H) Beta diversity using principal coordinate analysis (PCoA) of P10 and (I) P11 milk samples were created using the matrix of pair-wise distance between samples determined by Bray-Curtis dissimilarity using unique amplicon variants. (J) Microbial composition of taxonomy at the family level for saline and MIA dams at P10 (Saline: n = 6; MIA: n = 6) and P11 (Saline: n = 6; MIA: n = 7). (K) LEfSe biomarkers plots. These figures display taxa that are significantly more abundant in the milk of saline-treated dams (blue bars) versus MIA-treated dams (red bars) on P10 and P11. These taxa were identified based on their significant distributions (p < 0.05) and effect sizes (LDA score) larger than 2 for P10 (Saline: n = 6; MIA: n = 6) and for P11 (Saline: n = 7; MIA: n = 7). Among these taxa, MIA was associated with a higher level of *Pseudomonadaceae* (LDA score = 2.94, p=0.02) and *Christensenellaceae* (LDA score = 2.82, p = 0.02) on P10, and a greater expression of *Stenotrophomonas maltophilia* (LDA score= 3.45, p=0.025), *Ruminococcaceae* (LDA score = 3.35, p = 0.02) and *Lachnospiraceae* (LDA score= 3.85, p = 0.01) on P11. Data are expressed as mean ± SEM. *p < 0.05, ***p <0.001, MIA versus Saline; ^a^p < 0.05, ^aa^p < 0.01, main effect of time; ^b^p < 0.05, ^bb^p < 0.01, ^bbb^p < 0.001, main effect of MIA. LPS – lipopolysaccharide.

In order to examine the possibility that MIA challenge increased expression inflammatory cytokines that could be transferred to nursing offspring, we analyzed the concentration of milk IL-6. While there were no significant differences in IL-6 concentration between saline and MIA treated dams, there was a main effect of time on this measure (**Figure 2E)**. We also tested the idea that bound LPS could cross into maternal milk and be absorbed by offspring. LPS binds directly to LBP which triggers a downstream pro-inflammatory response and the release of acute phase proteins like IL-6 (46-47). LBP levels were measured in blood plasma and P10 milk samples. LPB levels were significantly greater in the blood plasma of MIA dams (**Figure 2F)**. In milk, LPB was absent in both MIA and saline dams. Additional data on maternal milk composition (i.e., protein, lactose, and IgA concentrations) can be found in **Supplementary Figure 2C-E**.

Microbiome sequencing revealed no main effect of treatment group on alpha (**Figure 2G**) or beta diversity in milk samples (**Figure 2H, I**). Milk samples collected on P11 demonstrated significantly greater alpha diversity along the Shannon index compared to samples collected on P10 (**Figure 2G**). Between treatment groups, LEfSE analysis identified nine differently abundant taxa on P10 and six significantly abundant taxa on P11 **(Figure 2J, K)**. These analyses revealed a greater abundance of *Pseudomonadaceae* and *Christensenellaceae* on P10, and a greater expression of *Stenotrophomonas maltophilia, Ruminococcaceae* and *Lachnospiraceae* on P11 in the milk from MIA dams. For additional data on the composition of the milk microbiome, see **Supplementary Figures 3-6**.

RNA-seq of the milk was performed to characterize the effects of mid-lactational MIA at the genome-wide level. A total of 829 genes were differently expressed on P10 between saline and MIA milk samples (**Figure 3A**). We analyzed clusters of gene networks that are involved in maternal milk nutrition, immune activation, and offspring brain development (48-55, **Figure 3B)**. Heatmap clustering revealed a general upregulation of genes related to nutrient transport and inflammation, while genes related to epigenetic modifications, parvalbumin development, and glucocorticoid signaling were largely downregulated by MIA on P10 (**Figure 3B**). In P11 samples (**Figure 4A, B)**, a total of 893 genes were differentially affected between the milk of saline and MIA dams. Heat clustering analysis of these samples demonstrated a general downregulation in genes responsible for inflammation, glucocorticoid signaling, epigenetic modifiers, and prolactin in MIA samples, while genes related to oxytocin, nutrient transport, triglycerides, and parvalbumin development were largely upregulated. For more information on the genes indicated in the heatmap clustering, see **Supplementary Tables 2 and 3**.

**Figure 3.**
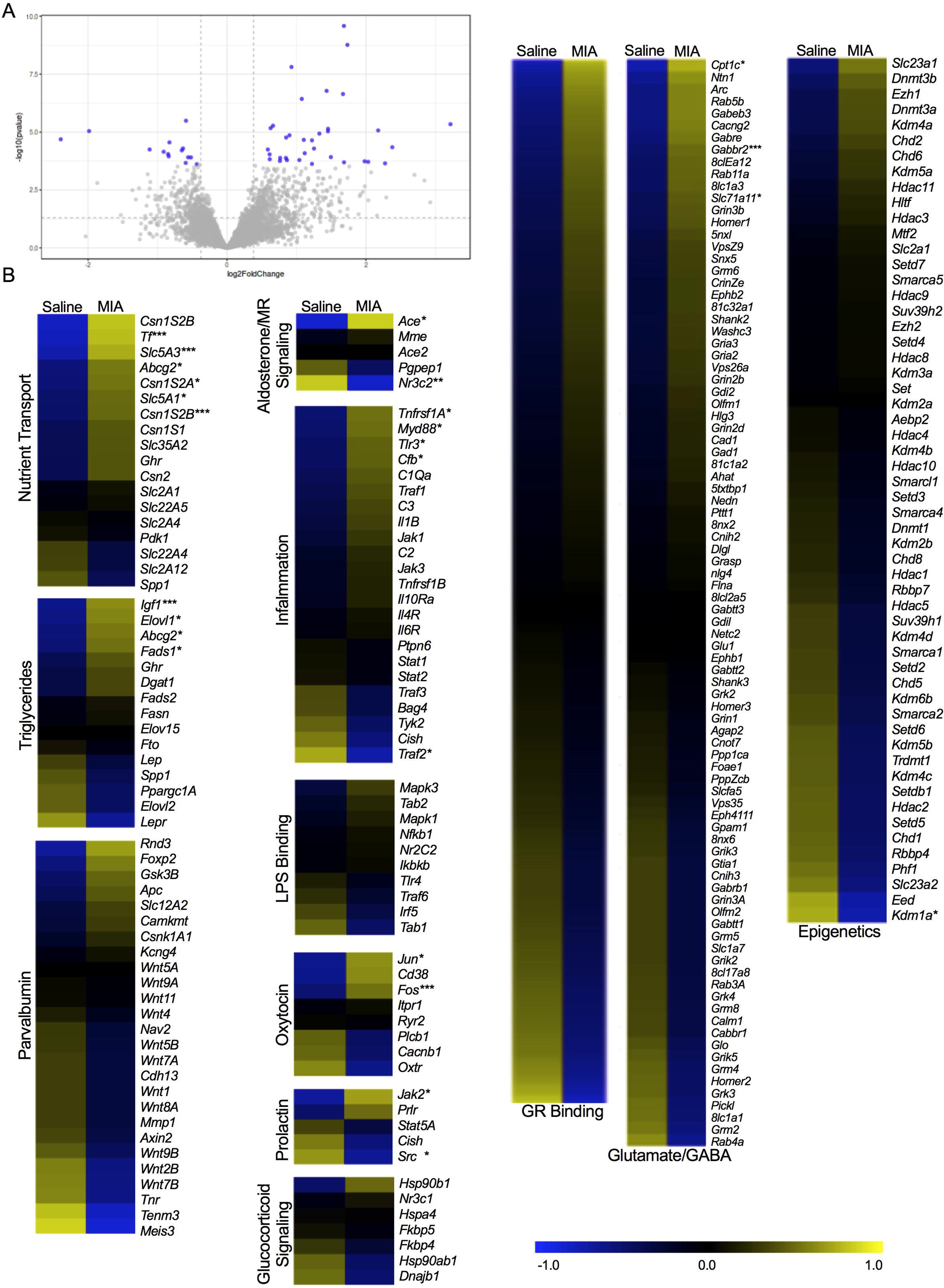
Transcriptomic analyses of milk samples obtained on P10 from rat mothers exposed to either saline or maternal immune activation (MIA). (A) Volcano plot depicting the distribution of 829 genes based on log2 fold change and -log10 p values. Grey dots are genes, and the dots highlighted in blue represent genes that displayed the highest magnitude of significance (padj<0.05, FC>1.3). (B) Heatmaps of differentially expressed genes related specifically to milk nutrient transport, triglyceride concentration, parvalbumin development, aldosterone/MR signaling, inflammation, LPS binding, oxytocin, prolactin, glucocorticoid signaling (events following glucocorticoids binding to the GR receptor), glucocorticoid binding, glutamate/GABA, and epigenetic modifiers. Expression is represented with the log2 transformation of counts recorded with a z-score based on the average across experimental groups. Data are expressed as *p <0.05 or **adjusted p <0.05, or ***FC>1.3, MIA versus Saline. GR-glucocorticoid receptor. MR-mineralocorticoid receptor.

**Figure 4.**
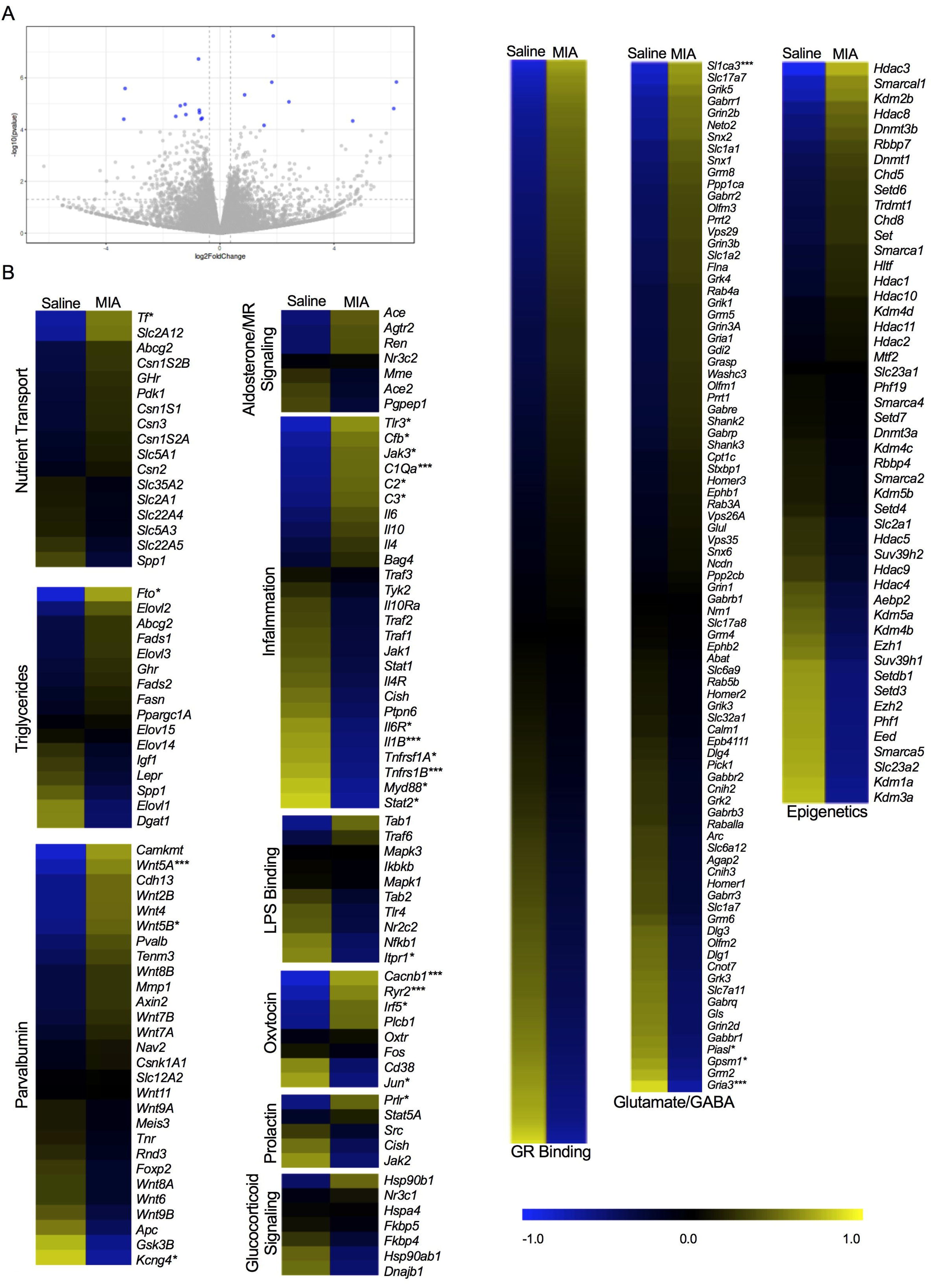
Transcriptomic analyses of milk samples obtained on P11 from rat mothers exposed to either saline or maternal immune activation (MIA). (A) Volcano plot depicting the distribution of 893 genes based on log2 fold change and -log10 p values. Grey dots are genes, and the dots highlighted in blue represent genes that displayed the highest magnitude of significance (padj<0.05, FC>1.3). (B) Heatmaps of differentially expressed genes related specifically to milk nutrient transport, triglyceride concentration, parvalbumin development, aldosterone/MR signaling, inflammation, LPS binding, oxytocin, prolactin, glucocorticoid signaling (events following glucocorticoids binding to the GR receptor), glucocorticoid binding, glutamate/GABA, and epigenetic modifiers. Expression is represented with the log2 transformation of counts recorded with a z-score based on the average across experimental groups. Data are expressed as *p <0.05 or **adjusted p <0.05, or ***FC>1.3, MIA versus Saline. GR-glucocorticoid receptor. MR-mineralocorticoid receptor.

### The combination of MIA challenge and nutritional deficit affected adolescent offspring physiology and behavior

Male, but not female MIA offspring were significantly heavier than saline controls on P43 (**Figure 5A**). See **Supplementary Figure 7** for additional data demonstrating the trajectory of weight gain in both offspring and dams across early development. Using body weight as a covariate, female offspring from MIA dams were found to have higher mechanical allodynia thresholds on the von Frey test compared to their saline counterparts (**Figure 5B**). This is suggestive of reduced sensitivity to tactile stimulation.

**Figure 5.**
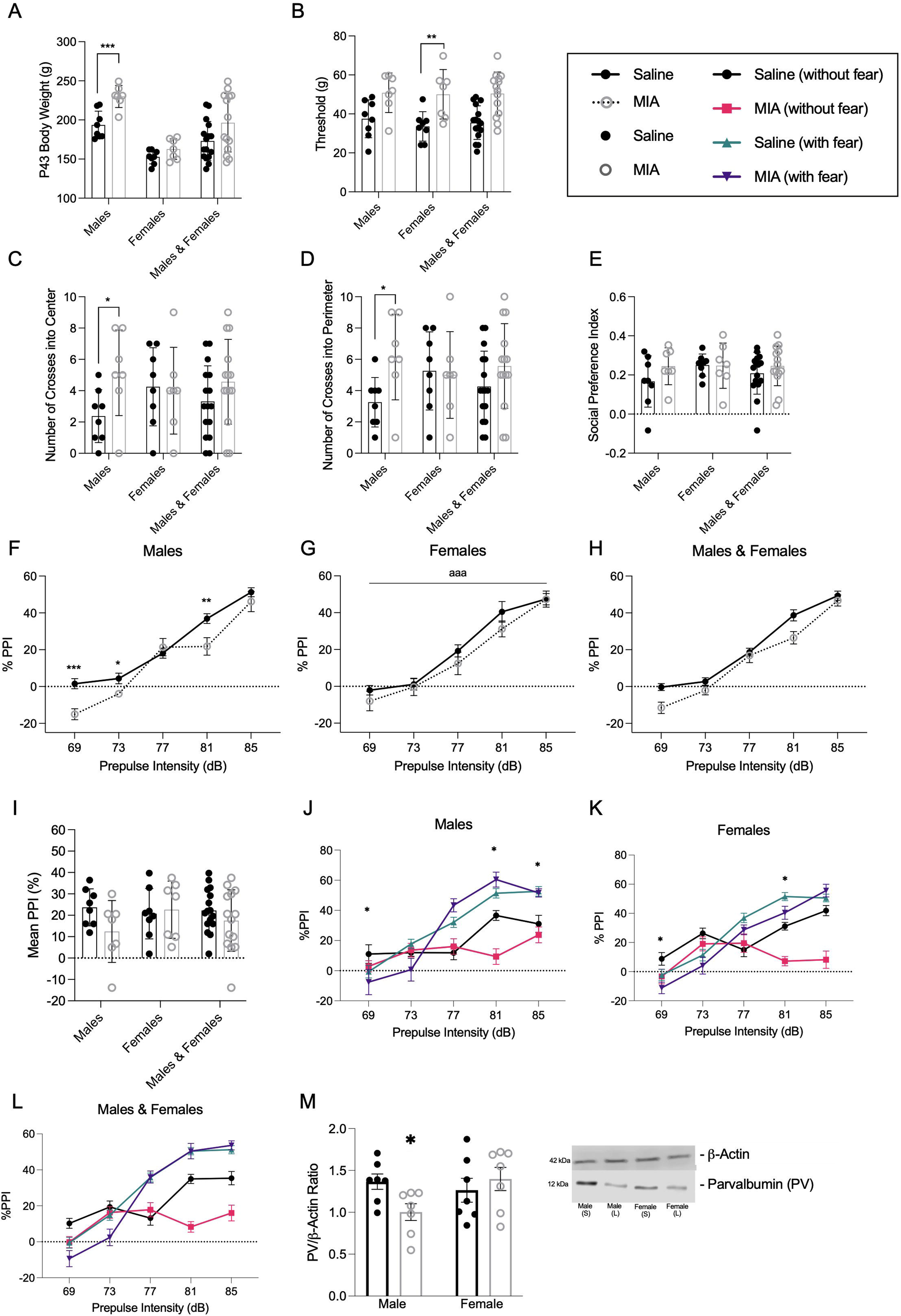
Adolescent offspring physiology and behavior following maternal immune activation (MIA) during the mid-lactational period. A main effect of MIA was observed in males for (A) postnatal day 43 body weights (grams; F(1,13) = 19.037, p = 0.001, *n*_*p*_^2^ = 0.594. (B) Mechanical paw withdrawal thresholds (grams) on the von Frey test evaluating mechanical allodynia were significantly greater in female offspring (F(1, 12) = 8.925, p =0.011, *n*_*p*_^2^ = 0.427). Male offspring made a greater number of crosses into the (C) center (F(1, 13) = 5.744, p = 0.032, *n*_*p*_^2^ = 0.306); and (D) perimeter (F(1, 13) = 6.513, p = 0.024, *n*_*p*_^2^ = 0.334) of an open field arena. There were no effects of MIA on (E) social preference index. Line plots displaying percent prepulse inhibition (PPI) as a function of increasing prepulse intensities for (F) males (significant MIA by time interaction F(4, 52) = 4.578, p = 0.003, *n*_*p*_^2^ = 0.260). Male offspring from MIA dams demonstrated significant reductions in PPI at the 69dB (F(1, 13) = 17.009, p = 0.001, *n*_*p*_^2^ = 0.743), 73dB (F(1, 13) = 5.927, p =0.030, *n*_*p*_^2^ = 0.581), and 81dB (F(1, 13) = 8.315, p = 0.013, *n*_*p*_^2^ = 0.633) intensities. (G) PPI performance for females (main effect of dB F(4, 52) = 80.357, p = 0.001, *n*_*p*_^2^ = 0.861), and (H) male and female PPI combined for display purposes. (I) The bar plot shows the mean percent prepulse inhibition across all five prepulse intensities. Conditioned fear % PPI for (J) males (main effects of MIA for 73dB (F(1, 13) = 6.787, p = 0.022, *n*_*p*_^2^ = 0.343), 81dB (F(1, 13) = 6.787, p = 0.022, *n*_*p*_^2^=0.343) and 85 dB (F(1, 13) = 13.124, p = 0.003, *n*_*p*_^2^=0.502)). Conditioned fear % PPI for (K), females (main effects of MIA for 69dB (F(1, 13) =12.969, p = 0.003, *n*_*p*_^2^=0.495) and 81dB (F(1, 13) = 25.117, p = 0.001, *n*_*p*_^2^=0.659)). (L) Males and females combined for display purposes. (M) Hippocampal parvalbumin (PV) and densitometric ratios for male and female offspring from saline (black) and MIA (grey)-treated dams. PV/β-actin ratios were significantly reduced in male MIA offspring (F(1, 13) =6.96, p = 0.022, *n*_*p*_^2^ = 0.367). Data are expressed as mean ± SEM; Saline: n = 8; MIA: n = 7 for the behavioral measures and Saline: n = 7; MIA = 7 for hippocampal PV. *p <0.05, **p<0.01, ***p <0.001, MIA versus Saline. ^a^p <0.05, ^aa^p<0.01, ^aaa^p <0.001, main effect of trial type.

Although there were no differences in distance traveled (cm), percent time spent in the center, duration of time (seconds) spent in the center, or perimeter of the open field arena (see **Supplementary Figure 8A-D**), male offspring of MIA exposed mothers displayed an increased number of crosses into both the center (**Figure 5C**) and perimeter (**Figure 5D**) of the arena. Social preference was not affected by the early life experience for either sex (**Figure 5E**).

Sensorimotor gating was evaluated using PPI of the acoustic startle reflex. A significant MIA by time interaction (**Figure 5F**) and a significant main effect of time (**Figure 5G, H**, collapsed across males and females for display purposes) confirmed that all groups demonstrated an increased % PPI as the intensity was raised from 69 to 85 dB. While females were unaffected, male offspring originating from MIA exposed mothers had attenuated PPI values compared to saline controls at the 69dB, 73dB, and 81dB intensities (**Figure 5F**). The mean percent prepulse inhibition across all five prepulse intensities was not affected as a function of maternal MIA exposure (**Figure 5I**).

With respect to conditioned fear, there was a main effect of MIA for % PPI during the trials without fear (**Figure 5J, K, L)**. % PPI for trials containing the conditioned stimulus was not significantly different between saline and MIA animals for either sex (**Supplementary Figure 8E**, collapsed across males and females for display purposes). One-way ANOVA revealed a main effect of trial type (**Supplementary Figure 8E**), which was expected given that the added element of fear increases % PPI (56-57). Conditioned fear did not lead to significant changes in total mean % PPI when collapsed across all dB, regardless of MIA or sex (**Supplementary Figure 8F**, data collapsed across males and females for display purposes), suggesting that fear modulated % PPI similarly regardless of sex or MIA exposure.

In addition to these behavioral effects, PV/β-actin ratios were significantly reduced in the hippocampus of male MIA offspring compared to saline dams (**Figure 5M**).

## Discussion

In the United States, postpartum bacterial infections develop in approximately 5-7% of mothers (58), and yet, little research is dedicated toward understanding how postnatal infection affects parental input signals like lactation and subsequent infant outcomes. We demonstrate that variations in milk quality may be part of a mechanism that programs offspring physiology and behavior following a transient stress exposure during the lactational period (see **Figure 6** for summary of proposed mechanisms). In the present study, acute MIA exposure on P10 was associated with sustained changes in both the nutritional content and microbial profile of maternal milk. Moreover, the MIA-induced changes in milk fat and corticosterone levels were accompanied by long-term physiological and behavioral alterations in offspring. For example, neonatal offspring from MIA-challenged dams spent less time huddling compared to controls. In adolescence, MIA offspring were heavier, demonstrated deficits in sensorimotor gating, threat detection, hippocampal PV expression, and had increased mechanical allodynia thresholds. These MIA-associated effects in offspring were unlikely due to an overexposure to inflammatory cytokines. Instead, offspring behavior and bodyweight differences likely reflected the unique expression of the nutritional elements and metabolically relevant bacterial taxa found in P10 and P11 milk samples. While most of the MIA-related literature has been dedicated to uncovering the physiological and behavioral effects on offspring gestationally exposed to MIA, fewer studies have focused on the impact of MIA exposure after birth. Importantly, the timing of MIA administration is critical to the translatability of these animal models to human studies, which have identified the second trimester as a sensitive window in cases of prenatal MIA exposure (2, 4). In rats, P10 corresponds to the perinatal period of development in humans (59), making our MIA model most relevant to maternal infections that occur soon after birth. Future studies are needed to extrapolate how MIA affects development when administered at later points during the lactational window.

**Figure 6.**
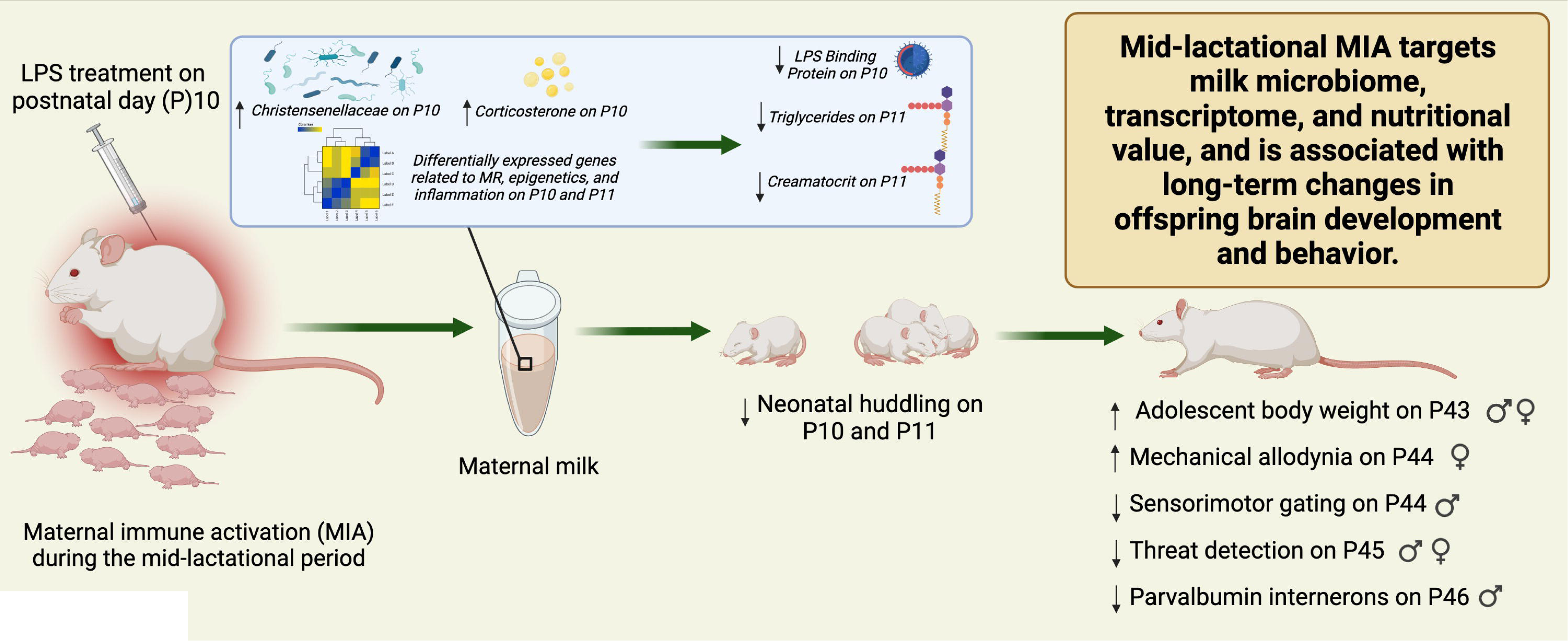
Summary of mechanisms affecting milk quality and offspring brain and behavior following maternal immune activation (MIA) during the mid-lactational period. Mid-lactational MIA experienced on postnatal day (P)10 is associated with changes in the milk transcriptome on both P10 and P11, in addition to increased milk corticosterone concentrations and microbial expression of *Christensenellaceae* on P10. In concert, these effects may lead to reduced LPS binding protein in milk on P10 and fat content on P11. We propose that this altered milk profile is one mechanism by which mid-lactational MIA contributes to the emergence of sex-specific behavioral and physiological changes across the early life of offspring. LPS – lipopolysaccharide; MR-mineralocorticoid receptor.

MIA was associated with multiple changes in the composition of maternal milk content. We observed a significant increase in milk corticosterone 2-hours post LPS treatment that subsided by P11. Milk corticosterone is expressed linearly to plasma corticosterone (60), validating our MIA model. Importantly, milk corticosterone can pass from a mother to her offspring where these glucocorticoids can remain active in the periphery and in the brain (61-62). Increased corticosterone exposure from breastmilk has previously contributed to HPA axis dysfunction as well as altered learning and memory abilities in rat pups of both sexes, independent of maternal care (63-64). In addition to its modulatory actions within the HPA axis, recent evidence has demonstrated the regulatory role of corticosterone in breastmilk where the presence of this hormone (the human equivalent of cortisol) can modulate nutritional elements including sodium, potassium (65-66) and fat (24). This evidence suggests that the temporary rise in corticosterone on P10 may have preceded downstream changes in milk nutritional quality. In line with this, MIA reduced % creamatocrit and milk triglycerides on P11. However, higher levels of glucocorticoids in milk alone are not a sufficient predictor of offspring bodyweight (24, 62), suggesting that the actions of corticosterone in neonates are likely interwoven with milk nutrition.

Microbiome sequencing did not reveal significant differences in alpha or beta diversity between saline and MIA treated dams. While general diversity was not affected, LEfSe analysis uncovered significant differences in the expression of certain taxa on P10 and P11. Specifically, MIA dams exhibited a higher abundance of bacterial families including *Pseudomonadaceae*, which was previously shown to be over expressed in the ceca of pregnant mice with low serum triglycerides (67). Further, milk from MIA dams also exhibited elevated levels of *Stenotrophomonas maltofilia*, which has the ability to degrade hydrocarbon chains and therefore could prevent triglyceride synthesis (68). Given that the microbes in breast milk directly colonize the infant gut (69-70), it is likely that these differently expressed taxa are being passed to the offspring to influence future metabolic processes. However, microbiome sequencing of the offspring duodenum was not performed, and this is an important limitation to the present study. Future experiments assessing the relationship between milk and the offspring microbiome will be critical to determine the contribution of these MIA-induced microbial differences to offspring development. Clinical research exploring potential therapeutic interventions, such as targeting the microbiome (71) which may rescue changes in milk composition and subsequent offspring developmental outcomes in contexts where postnatal infections are common, would highly benefit from these future investigations.

Differences in microbial and nutritional markers paralleled MIA-mediated changes found in the whole-transcriptome of the milk using RNA-seq. Notably, we observed a significant upregulation in aldosterone-promoting *Ace* and a downregulation of the mineralocorticoid receptor-coding *Nr3c2* in MIA milk on P10 (**Figure 3B, Supplementary Table 2**). These changes, coupled with the increase in P10 milk corticosterone, corroborated differences observed in milk nutrition given that aldosterone is another crucial mediator of breastmilk nutrition and ion exchange (65). These results were further supported by the differential genomic profiles between milk from MIA and saline dams for gene clusters related to triglycerides and nutrient transport. RNA-seq also revealed a general upregulation of genes involved in the inflammatory response in maternal milk on P10 in MIA-treated dams. Although LBP and IL-6 were equally regulated between saline and MIA dams, the possibility that other inflammatory cytokines may be active in milk and subsequently affect offspring development cannot be entirely ruled out. There was an unexpected significant upregulation of several genes within the inflammation pathway on P11 (**Figure 4B, Supplementary Table 3**), however this may be reflective of the simultaneous downregulation of epigenetic modifiers since cytokine release is often associated with aberrant epigenetic activity (72). While the function of breastmilk RNA once it is absorbed by offspring is still being unpacked, fluorescently tagged RNA-containing milk exosomes were shown to reach the offspring brain *in vivo*, and these exomes have the ability to influence cellular processes *in vitro* (73, 74). Our data identified several key genes related to brain development within maternal milk including Foxp2, Homer1, Tnr, and Gabbr2, necessitating a more focused investigation into how these genes may have been differently expressed in the offspring brain following mid-lactational MIA. Overall, these data demonstrate the extensive effects of MIA on maternal milk at the genome-wide level.

We used several behavioral measures to explore the role of lactational MIA exposure on offspring development. Neonatal ‘huddling’ is described as a form of social thermoregulation (75). One previous study found a reduction in nest seeking behavior in rat pups prenatally exposed to LPS (76). Undernourishment has also been shown to reduce neonatal huddling in rat pups (77), and huddling efficiency may be related to serum triglycerides in newborn rabbits (78). This evidence suggests that MIA-modulated huddling may emerge due to differences in the fat content of maternal milk. Disruptions in huddling can also arise following changes to maternal care (79-81). Although maternal care in the present study did not drastically differ between MIA and saline dams, we did observe differences in maternal self-directed care. Therefore, reduced pup huddling may not be exclusively promoted by MIA-associated changes in milk quality.

Prenatal MIA has been shown to modulate offspring performance on certain tasks, such as the open field, social preference, PPI, measures of mechanical allodynia, and fear conditioning paradigms (4, 36, 82-84). Here, we observed significant MIA induced reductions in % PPI that were exacerbated in male adult offspring while females demonstrated deficits in mechanical allodynia. MIA was also associated with a significant reduction in % PPI in trials without fear, following the foot shock and conditioned stimulus (CS) pairing. These findings suggest MIA contributes to an impairment in the offspring’s ability to discriminate between “safe” trials (without CS presentation) and “stressful” trials (with CS presentation). Early life stress has been shown to impair this form of contextual threat detection in adolescence, and this is especially true for females (85-86). These results, in addition to reduced neonatal huddling, point to generally impaired sensory processing and threat responses in the offspring of dams exposed to MIA during lactation. While we might expect these behavioral changes in our offspring to be maintained into adulthood, neonatal LPS exposure has been associated with altered behavior during specific developmental windows (e.g., adolescence) which can remit by later ages (e.g., adulthood; 87). Therefore, future studies should consider assessing both adolescents and adults in order to better understand the trajectory of these effects. The lactational period is an important time for neonates and here we underscore the value of maternal inputs (e.g., maternal milk quality and behavior) in offspring behavioral development.

We measured the expression of GABAergic PV in the offspring hippocampus since altered PV expression in this region is a common theme in prenatal MIA models and is associated with many MIA-associated behavioral and sensory processing changes (88-90). Further, poor nutrition in early life can prevent the development of PV in rat hippocampus (91) although this has only been explored in males. The emergence of neonatal huddling behavior is time-locked to the maturation of GABAergic signaling (92). Therefore, mid-lactational MIA may disrupt this early form of social behavior by attenuating the development of PV cells. Importantly, MIA dams showed significant increases in pup-directed licking and grooming behavior the day following MIA challenge. This heightened maternal care demonstrated by MIA dams on P11 may have been reflected by the upregulated expression of milk genes related to oxytocin on P11, but also may have acted as a compensatory mechanism to buffer the future development of more severe social deficits in adolescent offspring. This is plausible, given that tactile stimulation through the licking and grooming of pups is one form of maternal behavior that promotes brain development and modulates future offspring behavior (93-96). In adolescence, male MIA offspring demonstrated a reduced expression in hippocampal PV. In vivo inhibition of PV in the hippocampus has previously been shown to reduce PPI in male mice (97). Mechanistically, gestational exposure to MIA results in a more reactive HPA axis and altered hippocampal circuitry in male animals (36). In this study, we show that mid-lactational MIA may target similar mechanisms, as our male offspring demonstrated reduced PV expression and greater bodyweights, which may suggest greater stress levels compared to females. Together, these results demonstrate one neuronal target by which MIA during mid-lactation contributes to lasting changes in offspring brain development in males. More work is needed to discern the neuronal targets of mid-lactational MIA in females.

Although baseline maternal care was not assessed prior to milking procedures in the present study, future studies should consider how milking procedures may confound maternal care and affect offspring. Aside from a general pattern of decreased (but not significant) licking and grooming of pups on the afternoon of P10, we did not observe a drastic reduction in pup directed behaviors in MIA dams. The effects of LPS are highly dose dependent (98). While Vilela and colleagues (12) observed more dramatic deficits in maternal care following MIA, they implemented a dose of LPS 5-times higher than the dose used in the present study. Moreover, lactational exposure to LPS using the same dose as that used in the current study also resulted in improved maternal care (14). Importantly, LPS is a bacterial memetic that acts via the Toll-like receptor 4 pathway (99). Therefore, caution should be taken when interpreting the clinical relevance of our finding since work is needed in this field to establish if viral and other types of infections may affect lactation in a similar manner. Together, the results of our study suggest that stressors experienced during the neonatal period may interact with maternal care and lactation quality, affecting offspring physiology and behavior. Indeed, early life stress models affect both the parent(s) and the offspring; the dynamic impacts on each need to be considered as part of the mechanistic programming of the developmental trajectory.

## Conclusions

Maternal care and milk quality are important components for offspring development, brain health, and behavior. Here, we demonstrate that an acute maternal immune response during mid-lactation is sufficient to trigger changes in breastmilk quality and its microbiome profile, in addition to the physiology and behavior of adolescent offspring. These results contribute to the broader literature by suggesting that the effects of MIA on offspring development are not restricted to the prenatal window. Moreover, this study ties in the characteristics of lactation and nutrition as part of a mechanism contributing to the trajectory of offspring development following early life stress. Given that animal models of early life stress can impact both parents and their offspring, the quality of maternal milk should be considered among the variables investigated in future studies. Finally, this work underscores the importance of research focused on potential therapeutic interventions (e.g., nutritional lactation supplements) and necessitates a better representation of pregnant and nursing people to aim for increased equity and inclusivity in both basic and clinical research.

## Supporting information

Supplementary Table 1. MIA Reporting Checklist

## Funding and Disclosures

This project was funded by NIMH under Award Number R15MH114035 (to ACK) and the Massachusetts College of Pharmacy and Health Sciences (MCPHS) Center for Undergraduate Research (H.T). The authors wish to thank Mary Erickson, Sahith Kaki, Ada Cheng, Dr. Mattia Migliore, and Dr. Theresa Casey for their technical support in the early phases of this project. The authors would also like to thank the MCPHS Schools of Pharmacy and Arts & Sciences for their continual support, the Bioinformatic Resource Center at the Rockefeller University where the RNA-seq was performed, and the University of Massachusetts Boston where HD is a graduate student. Figure 6 made with BioRender.com and an earlier version of this manuscript was posted on the preprint server bioRvix. The content is solely the responsibility of the authors and does not necessarily represent the official views of any of the financial supporters.

## Author Contributions

H.D., S.G.C., H.T., and A.C.K., ran the experiments; H.D., S.G.C., J.M., & A.C.K. analyzed and interpreted the data; H.D. and A.C.K. wrote the manuscript; A.C.K., designed and supervised the study.

## Conflict of Interest

The authors declare that they have no known competing financial interests or personal relationships that could have appeared to influence the work reported in this paper.

## Supplementary Methods

**Supplementary Table 1**. Maternal immune activation model reporting guidelines checklist.

**Supplementary Table 2**. Differentially expressed genes in P10 milk samples from the selected pathways. Genes with “NA” padj values indicate that FDR was not calculable due to fewer normalized counts than the optimal threshold.

**Supplementary Table 3**. Differentially expressed genes in P11 milk samples within the targeted pathways.

**Supplementary Figure 1. Maternal care behaviors following maternal immune activation (MIA) during the mid-lactational period**. Total number of (A) pup retrievals, (B) passive nursing bouts, (C) nest building behaviors, and (D) maternal drinking bouts. Maternal care was evaluated in the afternoon and night of postnatal day (P) 10 and the morning of P11. Saline: n = 8, MIA: n = 7. Data are expressed as mean ± SEM. ^a^p < 0.05, main effect of time.

**Supplementary Figure 2**. Nutritional profile of milk following maternal immune activation (MIA) during the mid-lactational period. Maternal milk (A) fat concentration (g/L), (B) energy value (kcal/L), (C) protein concentration (mg/mL), (D) lactose concentration (ng/μL), and IgA concentration (mg/mL). Saline: n = 8; MIA: n = 7. Data are expressed as mean ± SEM. *p < 0.05, ***p <0.001, MIA versus Saline; ^a^p < 0.05, main effect of time.

**Supplementary Figure 3**. Cladogram of milk biomarkers associated with treatment group on P10 (determined by LEfSe). Diameter of the nodes indicates relative abundance of taxa for saline (green) and MIA (red) samples. Placement indicates the classification of taxa, where nodes decrease in rank the closer to the center of the diagram. (Saline: n = 6; MIA: n = 6).

**Supplementary Figure 4**. Taxonomy heatmap demonstrating the top fifty most abundant species identified in samples on P10. Treatment group is indicated by the colored bar at the top of the figure (red = saline, blue = MIA). Each row represents the abundance for each taxon, with the taxonomy ID shown on the right. Each column represents the abundance for each sample. (Saline: n = 6; MIA: n = 6).

**Supplementary Figure 5**. Cladogram of milk biomarkers associated with treatment group on P11 (determined by LEfSe). Diameter of the nodes indicates relative abundance of taxa for saline (green) and MIA (red) samples. Placement indicates the classification of taxa, where nodes decrease in rank the closer to the center of the diagram. (Saline: n = 7; MIA: n = 7).

**Supplementary Figure 6**. Taxonomy heatmap demonstrating the top fifty most abundant species identified in samples on P11. Treatment group is indicated by the colored bar at the top of the figure (red = saline, blue = MIA). Each row represents the abundance for each taxon, with the taxonomy ID shown on the right. Each column represents the abundance for each sample. (Saline: n = 7; MIA: n = 7).

**Supplementary Figure 7. Offspring and maternal body weights across early development following maternal immune activation (MIA) during the mid-lactational period**. A) Male body weights demonstrated a significant effect of time (F(2, 26)= 1078.59, p<0.001) and MIA (F(1,13)= 7618.25, p<0.001) while B) female offspring and C) dam body weights only differed significantly across time (females: F(2, 26)=1491.4, p<0.001; maternal body weight: F(1, 13)=21.64, p<0.001). P10 offspring body weights were obtained by taking the average weight from the weight of all pups of the same sex for each litter (Saline: n = 8; MIA n = 7). P21 and P43 body weights were obtained from one male and one female per litter (Saline: n = 8; MIA n = 7). Data are expressed as mean ± SEM. ***p < 0.001, main effect of MIA; ^aaa^p < 0.001, main effect of time.

**Supplementary Figure 8. Adolescent offspring behavior following maternal immune activation (MIA) during the mid-lactational period**. (A) Distance traveled (cm), (B) percent time spent in the center, and duration of time (seconds) spent in the (C) center and (D) perimeter of an open field arena. (E) Conditioned fear significantly increased %PPI regardless of MIA or sex (F(4, 52)= 8.19, p=0.008, *n*_*p*_^*2*^=0.226). (F) %PPI for trials primed with conditioned fear collapsed across males and females for display purposes. Saline: n = 8; MIA n = 7. Data are expressed as mean ± SEM. Data are expressed as mean ± SEM, *p<0.05.

## Notes

### Competing Interest Statement

The authors have declared no competing interest.

### Summary of Updates

Figure 5

