## Supplementary Methods for "Got Milk? Maternal immune activation during the mid-lactational period affects nutritional milk quality and adolescent offspring sensory processing in male and female rats"

### Supplementary Methods

#### Animals and Housing

Male and female Sprague-Dawley rats (Charles River, Wilmington, MA) were housed in same-sex pairs and maintained at 20°C on a 12 h light/dark cycle (0700-1900 light) in standard sized laboratory cages (27×48×22 cm). Food and water were available *ad libitum* throughout the study. During breeding, animals were placed into larger cages (51 x 41 x 22 cm) and bred using a 1 male:2 female design. Pregnancy was confirmed by continued weight gain and visible teats during the later phase of gestation. Approximately two days prior to parturition, females were individually housed in fresh clean standard sized cages. Day of birth was designated as postnatal day (P)0 and litters were standardized to 10 pups per litter on P1. Wherever possible, litters were balanced with equal numbers of male and female pups. Animals stayed in this condition until they were weaned into same-sex pairs on P22, again using standard sized laboratory cages. All cages maintained a tube, a Nylabone chew toy®, and Nestlets® throughout the study, except during the two-day period prior to parturition through until P14 when toys were removed to protect pups from injury. Additional Nestlets® were available to litters during this time. One male and one female offspring from each litter were evaluated on behavioral tasks beginning on P43. The behavioral monitoring equipment was cleaned thoroughly with the disinfectant cleaner Quatriside TB (Pharmacal Research Laboratories, Inc) between each animal and test. The inter-rater reliability between blinded scorers was found to be greater than r=0.80 for each observational measure reported in this study.

#### Maternal Immune Activation (MIA)

On the morning of P10, dams were removed from their litter and placed into a fresh clean cage located in a separate procedure room. To model MIA during the lactational period, dams were weighed and intraperitoneally (i.p.) administered either 100 µg/kg of the inflammatory endotoxin, lipopolysaccharide (LPS; Escherichia coli, serotype 026:B6; L-3755, Sigma, St. Louis, MO, USA; n = 7) or pyrogen-free saline (saline; n = 8) at the start of the separation period. Animals remained in this cage for 2 hours until milking. The purpose of the separation period was to allow for maternal milk accumulation so samples could be collected at the height of the inflammatory response. In their regular holding room, pups were placed into a smaller clean cage positioned on top of a heating pad to maintain their body temperature. Pups were weighed immediately prior to being returned to their dams and again 2 and 24 hours later, alongside the inspection of milk bands, to monitor their health. Twenty-four hours following the P10 inflammatory challenge, milk collection procedures were repeated to evaluate sustained changes in milk quality. Additional details can be found in the methodological reporting table from Kentner et al. (2019), provided as Supplementary Table S1.

#### Milk Sample Collection and Composition

Milk was collected from each dam on P10 and P11 following a modified procedure from Paul et al. (2015). Immediately following each 2-hour separation period, dams were lightly anesthetized with isoflurane in O2 followed by the administration of 0.2 mL of oxytocin (20 USP/mL i.p.). The pharmacokinetic profile of isoflurane suggests that the drug is not absorbed by offspring and that breastfeeding can resume immediately after anesthesia (Drugs and Lactation Database, 2020; Lee & Rubin, 1993). Similarly, oxytocin is not expected to affect offspring given its short plasma half-life of 1-6 minutes which is reduced even further during lactation (Par

Pharmaceutical Inc, 2020). Teats were prepared by moistening the collection areas with distilled water. Milk was obtained from each dam by gently squeezing the base of the teat and manually expelling the milk for collection. At the time of milk collection, ~20 ul of sample was collected into a microhematocrit tube, which was then sealed and placed into a hematocrit spinner (2 minutes at 13, 700 g; StatSpin CritSpin Microhematocrit Centrifuge, Beckman Coulter, Inc). Measurements were taken to calculate percent (%) creamatocrit by evaluating the sample separation into each of the cream and clear layers by following the procedures outlined in Paul et al. (2015). The remaining milk that was collected (~500 ul per animal) was aliquoted across microcentrifuge tubes and stored at - 80°C until further processing. Collection time took about 10-25 minutes per rat, and dams were reunited with their litters in their home cage as soon as they fully awoke from anesthesia.

Milk samples were homogenized by overnight rotation on a Mini Tube Rotator (Fisher Scientific Cat. #88861051) at 4°C before analysis. A lactose assay kit (Sigma-Aldrich Cat. #MAK017) was used to measure lactose content in milk diluted 1:500 with lactose assay buffer (as outlined by Chen et al., 2017 and DeRosa et al., 2022). Triglycerides were evaluated using a colorimetric assay kit (Abcam, Cat. #ab65336) at 1:1000 dilution and the Pierce™ BCA Protein Assay Kit (Cat. #23227) was run at a 1:50 dilution to measure protein levels in milk (DeRosa et al., 2022). The small sample assay protocol of the corticosterone ELISA kit (#ADI-900–097, Enzo Life Sciences, Farmingdale, NY) was followed, as recommended by the manufacturer, using a 1:40 dilution. Milk levels of immunoglobulin (Ig) A were assessed with an assay kit (Bethyl Laboratories, Cat. # E111-102) using samples diluted to 1:1000 (DeRosa et al., 2022) while interleukin (IL)-6 (Thermo Fisher Scientific, Cat. #BMS625) was evaluated using samples diluted 1:2. Undiluted milk samples and blood plasma (diluted 1:2) were processed using the LPS binding protein ELISA (Abcam, Cat. #ab269542). Manufacturer's instructions were followed for all ELISA kits (n = 7).

**Microbiome Sequencing**

P10 (Saline: n = 6; MIA: n=6) and P11 (n = 6-7) milk samples underwent microbiome sequencing. DNA extraction was performed using the ZymoBIOMICS ®-96 MagBead DNA Kit (Zymo Research, Irvine, CA) and 16S targeted sequencing was completed using the Quick16S™ NGS Library Prep Kit (Zymo Research, Irvine, CA) and V3-V4 16S primers (Zymo Research, Irvine, CA). The sequencing library was prepared using real-time PCR. Final PCR products were quantified with qPCR fluorescence readings and pooled together based on equal molarity. The final pooled library was cleaned with the Select-a-Size DNA Clean & Concentrator™ (Zymo Research, Irvine, CA) which was then quantified with TapeStation® (Agilent Technologies, Santa Clara, CA) and Qubit® (Thermo Fisher Scientific, Waltham, WA). The final library was sequenced on Illumina® MiSeq™ with a v3 reagent kit (600 cycles) and sequencing was performed with 10% PhiX spike-in. Unique amplicon sequence variants were inferred from raw reads using the DADA2 pipeline (Callahan et al., 2016). Potential sequencing errors and chimeric sequences were also removed with the DADA2 pipeline. Taxonomy assignment was performed using Uclust from Qiime (v.1.9.1) and referenced with the Zymo Research Database (Zymo Research, Irvine, CA).

**RNA Sequencing**

Following homogenization, total RNA (n=4-5 per group) was isolated from a subset of milk samples following the protocol described by Chen et al. (2017) and DeRosa et al. (2022). The miRNeasy Mini Kit (QIAGEN, Cat. #217004) was used to isolate total RNA following the manufacturer's directions. Isolated RNA was quantified utilizing a NanoDrop 2000 spectrophotometer (ThermoFisher Scientific). RNA samples were stored at -80°C until RNA-sequencing (RNA-seq) which was performed at the New York Genomic Center, Libraries were processed through Tapestation to determine the concentration of DNA and fragment size. 100-bp reads were obtained using an Illumina NovaSeq 6000. Samples were read at a sequencing depth of approximately 50 million reads. A $p < 0.05$, Benjamini–Hochberg false discovery rate corrected (FDR) and fold change (FC) > 1.3, were identified using DESeq2 package and used to determine differentially expressed genes (Love et al., 2016). Heatmaps of differentially expressed genes were curated using normalized counts and were graphed with the MultiExperiment Viewer (National Library of Medicine, USA). Gene ontology (GO) analysis utilizing genes with $p < 0.05$ and FC > 1.3 was performed using the Database for Annotation, Visualization and Integrated Discovery functional annotation cluster tool (https://david.ncifcrf.gov/).

**Neonatal Huddling Behavior**

Two-hours after being reunited with their dams on P10 and P11, male and female offspring were quickly weighed and evaluated on their huddling behavior (Full litters evaluated togehter; Saline: n = 7; MIA: n = 7). This paradigm was adapted from Naskar and colleagues (2019). Briefly, litters were uniformly placed along the perimeter of a 40 cm x 40 cm area and video recorded for 10 minutes. The average number of pup clusters (2 or more pups in physical contact) was determined by extracting one video frame every 30 seconds and averaging the number of clusters observed across 20 frames. The average time spent together was determined by calculating the average time (in seconds) each pup spent in a clump for the entire 10-minute video.

**Maternal Behavior**

To evaluate whether lactational MIA exposure affected maternal behavior, we monitored passive maternal care between P10 and P11 (Saline: n = 8; MIA: n = 7). The first observation took place on the afternoon of P10 (15:00-17:00 hrs), following the inflammatory challenge. Maternal care observations took place again at 20:00-22:00 hrs, and then at 07:30-9:30 hrs on P11. Following the procedures of Strzelewicz et al (2021; 2019), each session consisted of six observations and a total composite score was calculated for each of the morning, afternoon, and evening time points. Dams were evaluated for 1-minute intervals per observation, with at least 5 minutes of no observations occurring between each of the 1-minute bins. Maternal care observations recorded included the frequency of pup-directed behaviors (i.e., dam licking/grooming pup, active/high crouch nursing, passive/low crouch nursing, pup retrieval), self-directed behaviors (i.e., dam eating/drinking, dam self-grooming), and nest building/digging behavior. Total time the dam spent on nest (seconds) was also recorded.

**Adolescent Open Field and Social Behavior**

On P43, male and female offspring were habituated to an open field arena for five-minutes (40 cm × 40 cm × 28 cm; Duque-Wilckens et al., 2020; Williams et al., 2020; Saline: n = 8; MIA: n = 7). An automated behavioral monitoring software program (Cleversys TopScan, Reston, VA) was used to evaluate animals on their duration spent (seconds) and frequency of crosses made in each of the center and perimeter of the arena. Percent time spent in the center of the arena and total distance traveled (cm) were also evaluated.

Immediately following the open field habituation period, two clean wire cups were placed on opposite ends of the arena to evaluate social preference (adapted from Crawley, 2007). One cup contained a novel object, and the other cup contained a novel rat of the same sex, age, and strain. Placement of the cups were counterbalanced between trials. Using a manual behavioral monitoring program (ODLog™ 2.0, http://www.macropodsoftware.com/), active investigation was recorded when an experimental rat directed its nose within 2 cm of a wire cup, or it was touching the cup. A social preference index was calculated by the formula ([time spent with the rat] / [time spent with the inanimate object + time spent with the rat]) - 0.5 (Scarborough et al., 2020).

**Mechanical Allodynia**

The next day, each animal was acclimatized for 30 minutes to an acrylic cage, with a wire grid floor. Mechanical allodynia was assessed using a pressure-meter which consisted of a hand-held force transducer fitted with a polypropylene rigid tip (Electronic von Frey Aesthesiometer, IITC, Inc, Life Science Instruments, Woodland Hills, CA, USA). The polypropylene tip was applied vertically to the central area of the animal's left hind paw with increasing force. The trial ended when the rat withdrew their paw from the tip, at which point the intensity of the stimulus was automatically recorded by the electronic pressure-meter. The average of four test trials was calculated as the mechanical withdrawal threshold (grams; Yan & Kentner, 2017).

**Prepulse Inhibition of the Acoustic Startle Reflex**

Three hours following the evaluation of mechanical allodynia, animals were placed into startle chambers and evaluated on prepulse inhibition (PPI) of the acoustic startle reflex (San Diego Instruments, San Diego, CA, USA). Sessions each consisted of pulse-alone, prepulseplus-pulse and prepulse-alone trials, as well as no-stimulus trials in which only a background noise of 65 dB was presented Giovanoli et al., (2013). The acoustic startle software administered one 40-ms pulse of white noise (120 dB) in combination with one of five different prepulses. The prepulses were made up of a 20-ms burst of white noise at five different intensities (69, 73, 77, 81, 85 dB), corresponding to 4, 8, 12, 16, and 20 dB above the background noise). After a startle habituation phase, each trial stimulus was pseudorandomly presented 12 times. The average interval between successive trials (ITI) was 15 ± 5 sec. Sessions terminated with 6 consecutive pulse-alone trials. PPI was calculated as percent inhibition of the startle response obtained in the prepulse-plus-trials compared to pulse-alone trials: [1-(mean reactivity on prepulse-plus-pulse trials / mean reactivity on pulse-alone trials) × 1/100] and expressed as % PPI for each animal at each of the five possible prepulse intensities.

**Conditioned Fear**

Fear conditioning has been shown to modulate PPI of the acoustic startle reflex (Ishii et al., 2010; Balogh et al., 2002). At the conclusion of the PPI session, animals were allowed to rest for 2 hours before commencing Day 1 of fear conditioning trials. On Day 1, animals received 10 trials of a light stimulus (conditioned stimulus; CS) paired with a 0.6 mA foot shock (unconditioned stimulus; US). The next day (Day 2), animals completed the PPI task for a second time, with half of the pseudorandom trials beginning with the presentation of the CS in order to assess how the element of fear influences PPI of the acoustic startle reflex. Percent PPI was calculated (described above) for trials with and without the CS.

**Tissue Collection**

On P46, a mixture of ketamine/xylazine (150 mg/kg, i.p./50 mg/kg, i.p.) was used to anesthetize animals. Whole hippocampus was dissected, frozen on dry ice and stored at −80°C for future processing. Maternal blood was collected at weaning in EDTA coated tubes following a cardiac puncture and spun at 1,000g for 10 minutes to obtain plasma.

**Western Blotting**

Offspring hippocampus was homogenized in a radioimmunoprecipitation assay buffer (RIPA) with protease inhibitor added (Thermo Fisher Scientific, Cat. #87786). Protein values were determined using the BCA assay (Pierce™, Cat. #23227). Samples were mixed with an equal volume of 2x Laemmli sample buffer (Bio Rad Laboratories Cat. #1610737) and denatured at 100°C for 5 minutes. 20µg of protein was loaded into each well of MiniProtean® gels (Bio Rad Laboratories, Cat. #4568101) and transferred onto nitrocellulose membranes (Bio Rad Laboratories, Cat. #1620147). Membranes were blocked in 5% nonfat milk with TBS + 0.05% Tween 20 (TBST) for 1 hour at room temperature, washed with TBST, and incubated in a 1:1000 dilution of parvalbumin (PV) antibody (RnD Systems, Cat. # AF5058) plus TBS overnight at 4°C. The next morning, membranes were washed with TBST and incubated in an HRP-conjugated secondary antibody (1:1000, RnD Systems, Cat., #HAF016) made in 1% nonfat milk with TBS for 1 hour at room temperature. Membranes were then washed with TBST and exposed to a chemiluminescent substrate (Thermo Fisher Scientific, Cat. #34580) for 5 minutes prior to being scanned. After imaging, membranes were stripped (Thermo Fisher, Cat. # 21062) for 15 minutes at 37°C, blocked in 5% nonfat milk with TBST for 1 hour at room temperature, washed, and incubated in beta actin primary antibody (1:1000, Thermo Fisher Scientific, Cat. #MA515739) for 1 hour at room temperature. Membranes were exposed again to the chemiluminescence substrate for 5 minutes and imaged. Densitometry was used to obtain a ratio of PV/β-actin in order to quantify differences between groups.

**Data Availability**

RNA-seq data have been deposited to GEO (GSE202525).
