## Supplementary Figures for "Got Milk? Maternal immune activation during the mid-lactational period affects nutritional milk quality and adolescent offspring sensory processing in male and female rats"

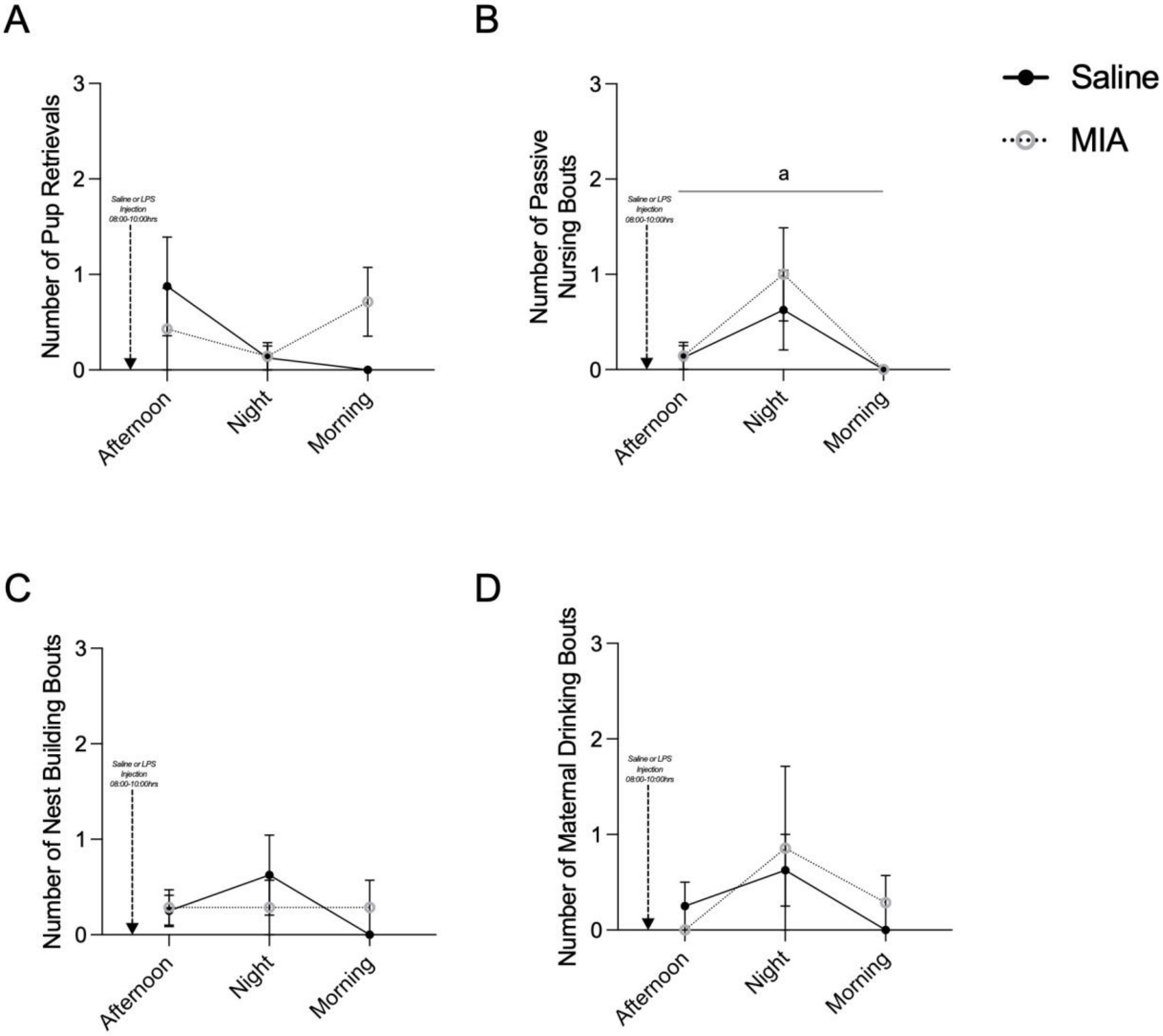


**Supplementary Figure 1. Maternal care behaviors following maternal immune activation (MIA) during the mid-lactational period.** Total number of (A) pup retrievals, (B) passive nursing bouts, (C) nest building behaviors, and (D) maternal drinking bouts. Maternal care was evaluated in the afternoon and night of postnatal day (P) 10 and the morning of P11. Saline: n = 8, MIA: n = 7. Data are expressed as mean ± SEM. [a]$p < 0.05$, main effect of time.

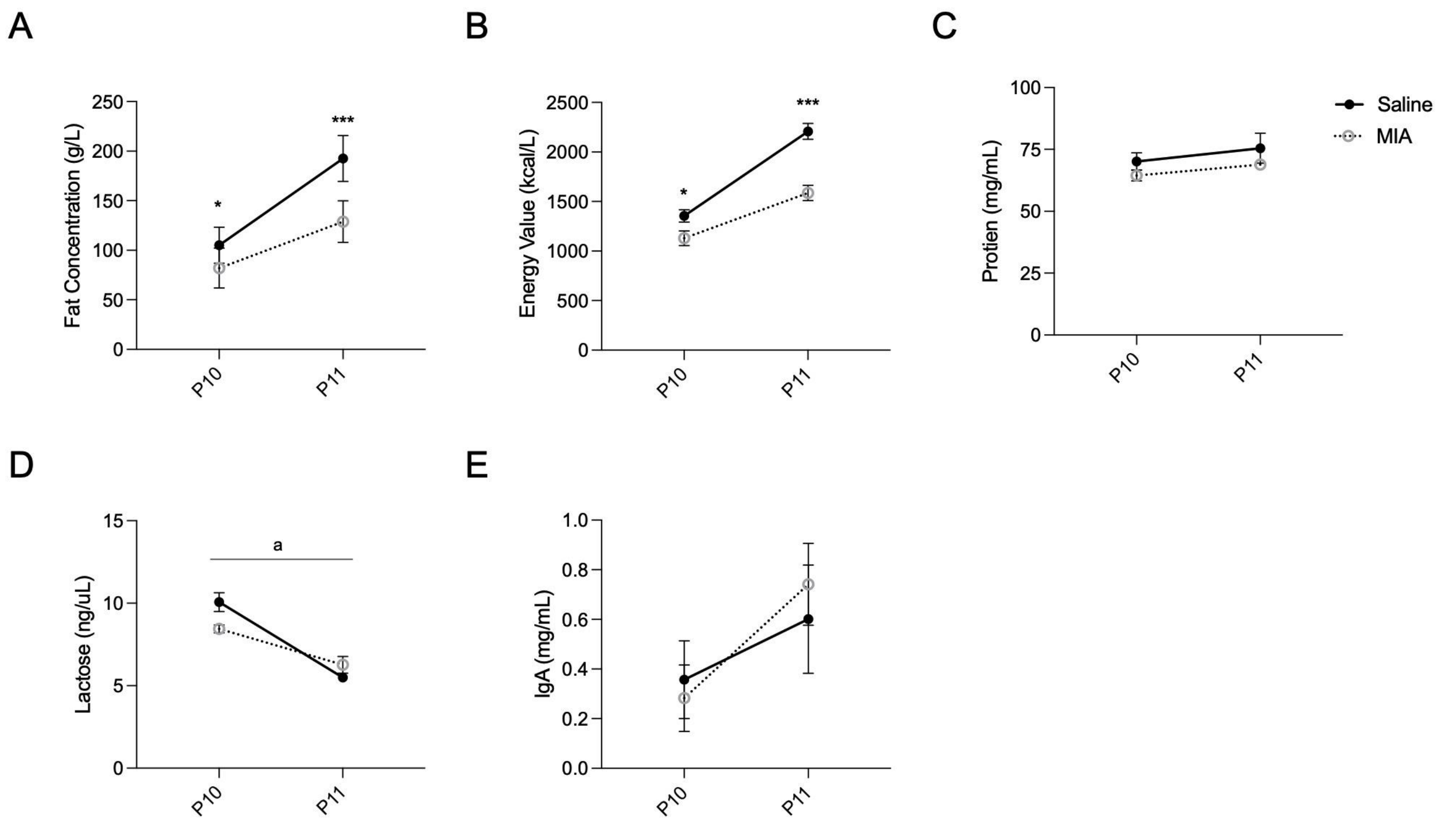


**Supplementary Figure 2.** Nutritional profile of milk following maternal immune activation (MIA) during the mid-lactational period. Maternal milk (A) fat concentration (g/L), (B) energy value (kcal/L), (C) protein concentration (mg/mL), (D) lactose concentration (ng/µL), and IgA concentration (mg/mL). Saline: n = 8; MIA: n = 7. Data are expressed as mean ± SEM. *$p < 0.05$, ***$p < 0.001$, MIA versus Saline; [a]$p < 0.05$, main effect of time.

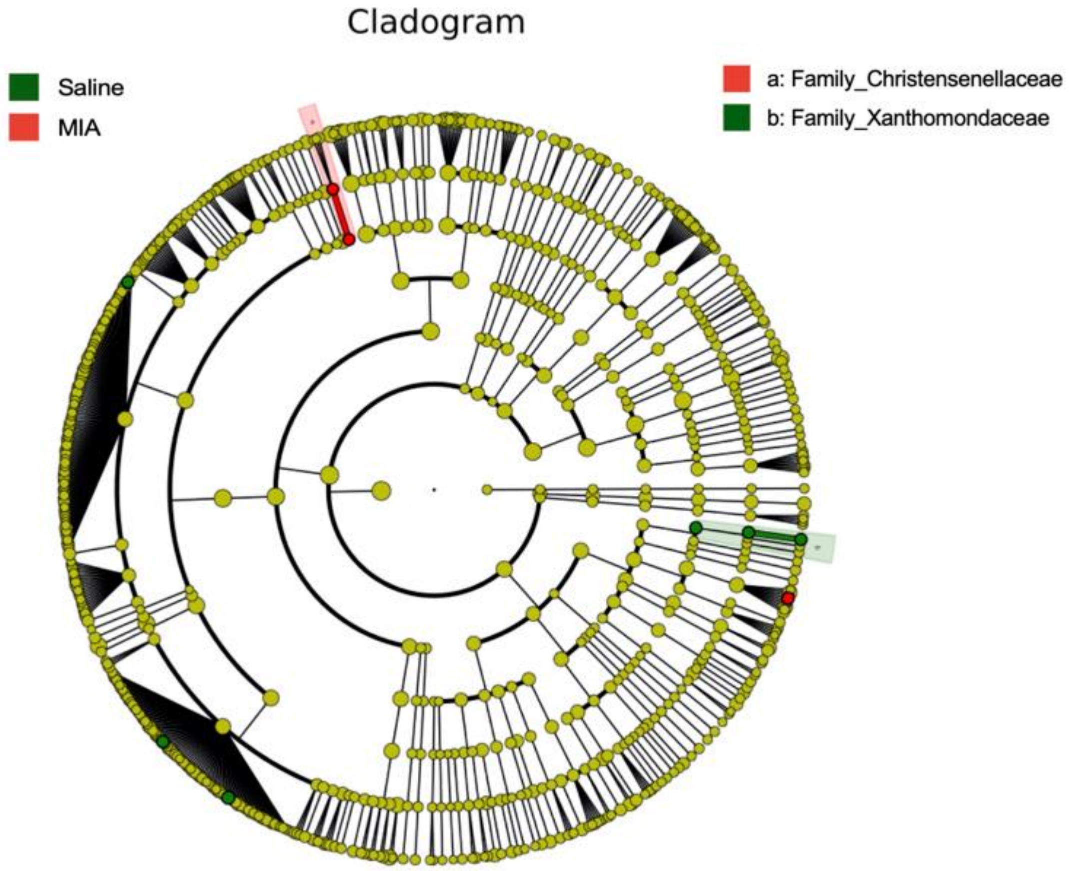


**Supplementary Figure 3.** Cladogram of milk biomarkers associated with treatment group on P10 (determined by LEfSe). Diameter of the nodes indicates relative abundance of taxa for saline (green) and MIA (red) samples. Placement indicates the classification of taxa, where nodes decrease in rank the closer to the center of the diagram. (Saline: n = 6; MIA: n = 6).

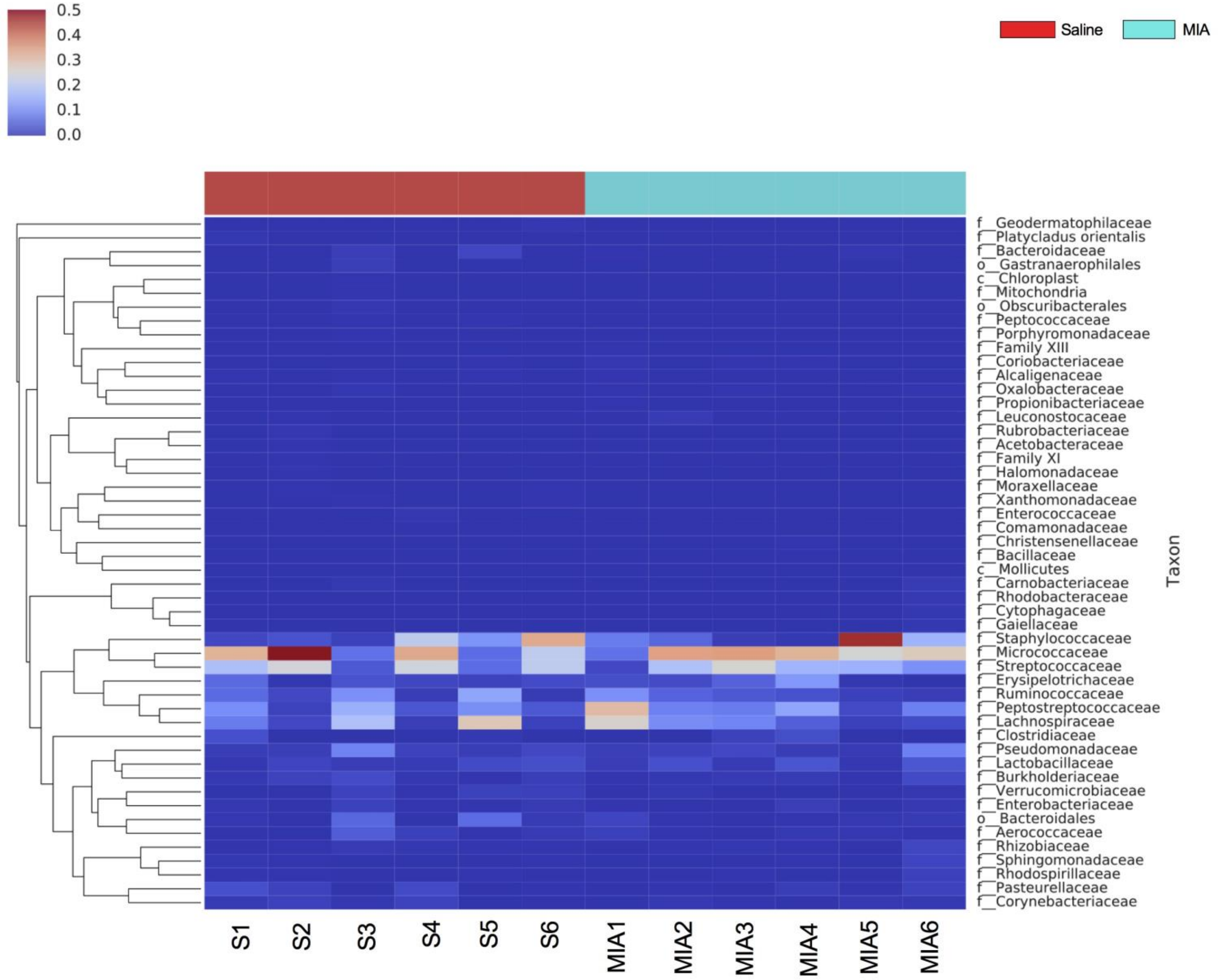


**Supplementary Figure 4.** Taxonomy heatmap demonstrating the top fifty most abundant species identified in samples on P10. Treatment group is indicated by the colored bar at the top of the figure (red = saline, blue = MIA). Each row represents the abundance for each taxon, with the taxonomy ID shown on the right. Each column represents the abundance for each sample. (Saline: n = 6; MIA: n = 6).

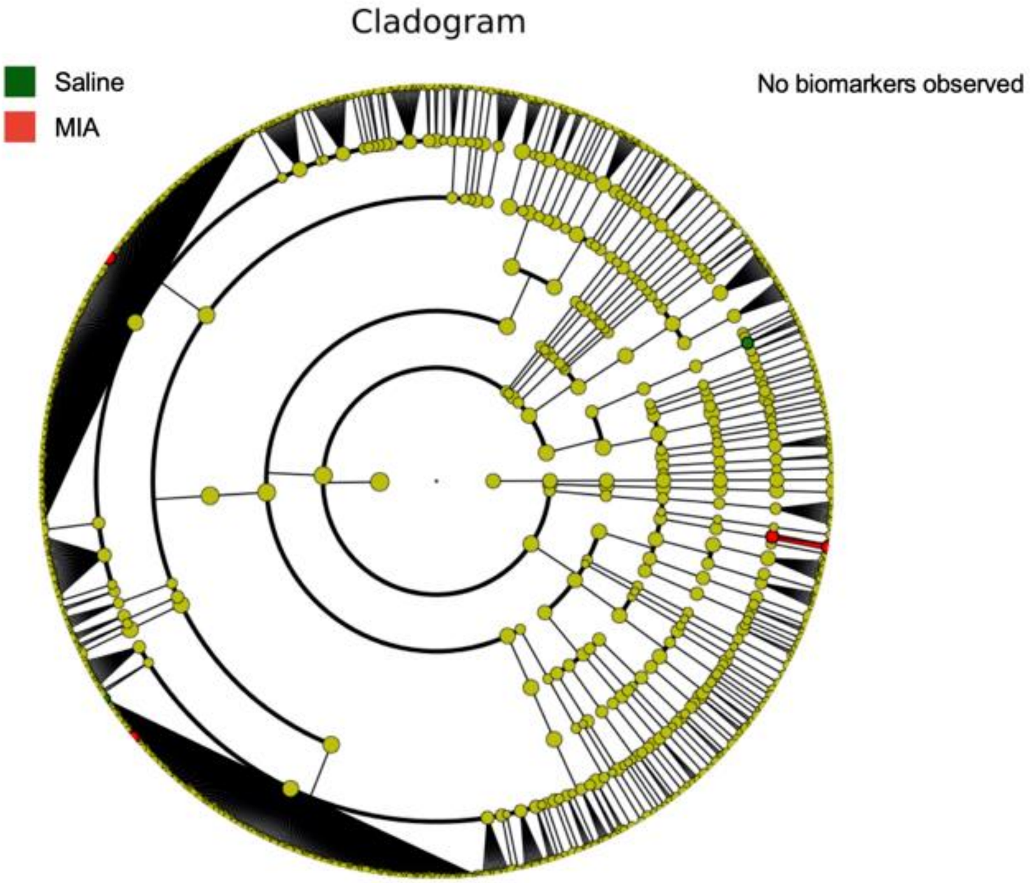


**Supplementary Figure 5.** Cladogram of milk biomarkers associated with treatment group on P11 (determined by LEfSe). Diameter of the nodes indicates relative abundance of taxa for saline (green) and MIA (red) samples. Placement indicates the classification of taxa, where nodes decrease in rank the closer to the center of the diagram. (Saline: n = 7; MIA: n = 7).

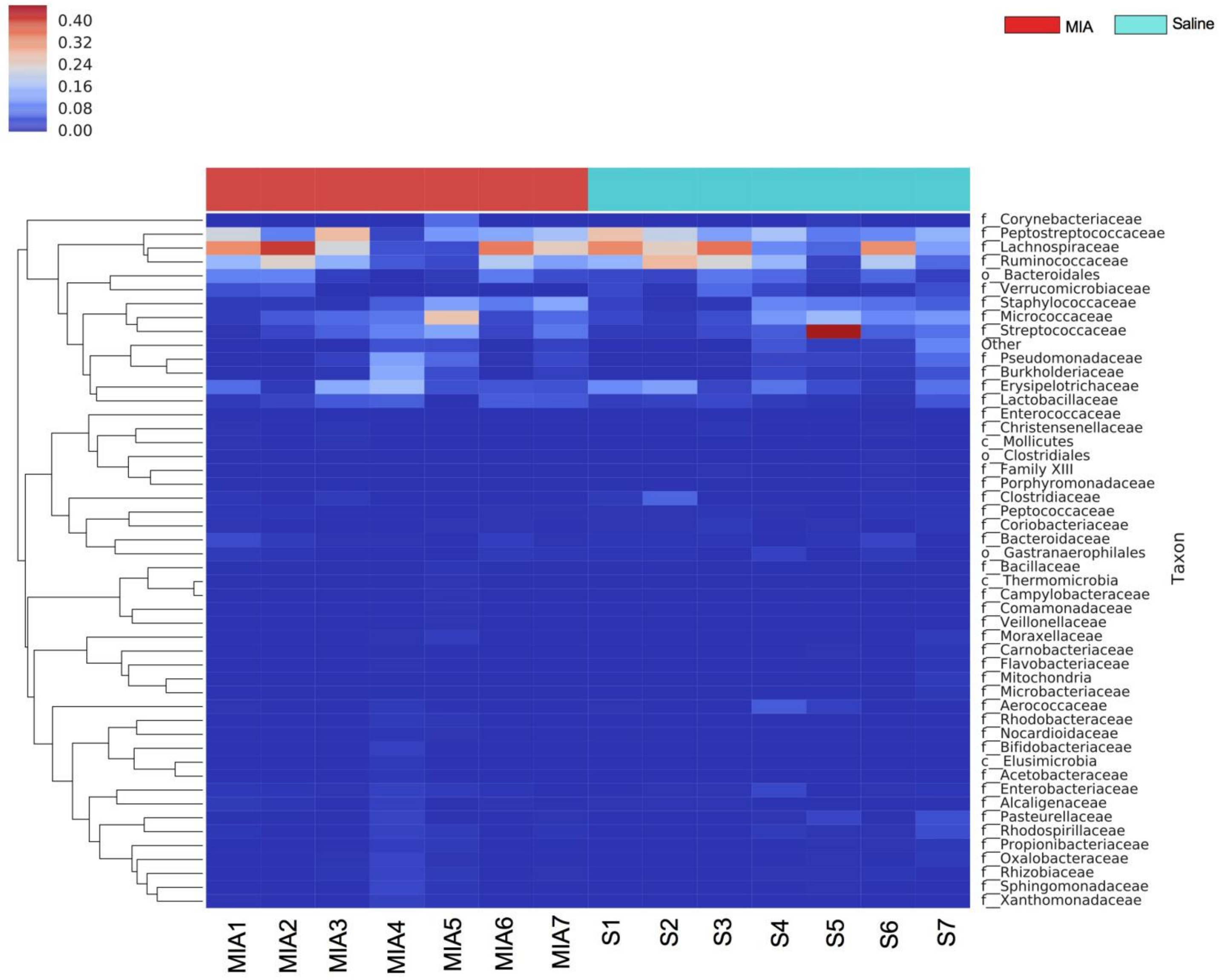


**Supplementary Figure 6.** Taxonomy heatmap demonstrating the top fifty most abundant species identified in samples on P11. Treatment group is indicated by the colored bar at the top of the figure (red = saline, blue = MIA). Each row represents the abundance for each taxon, with the taxonomy ID shown on the right. Each column represents the abundance for each sample. (Saline: n = 7; MIA: n = 7).

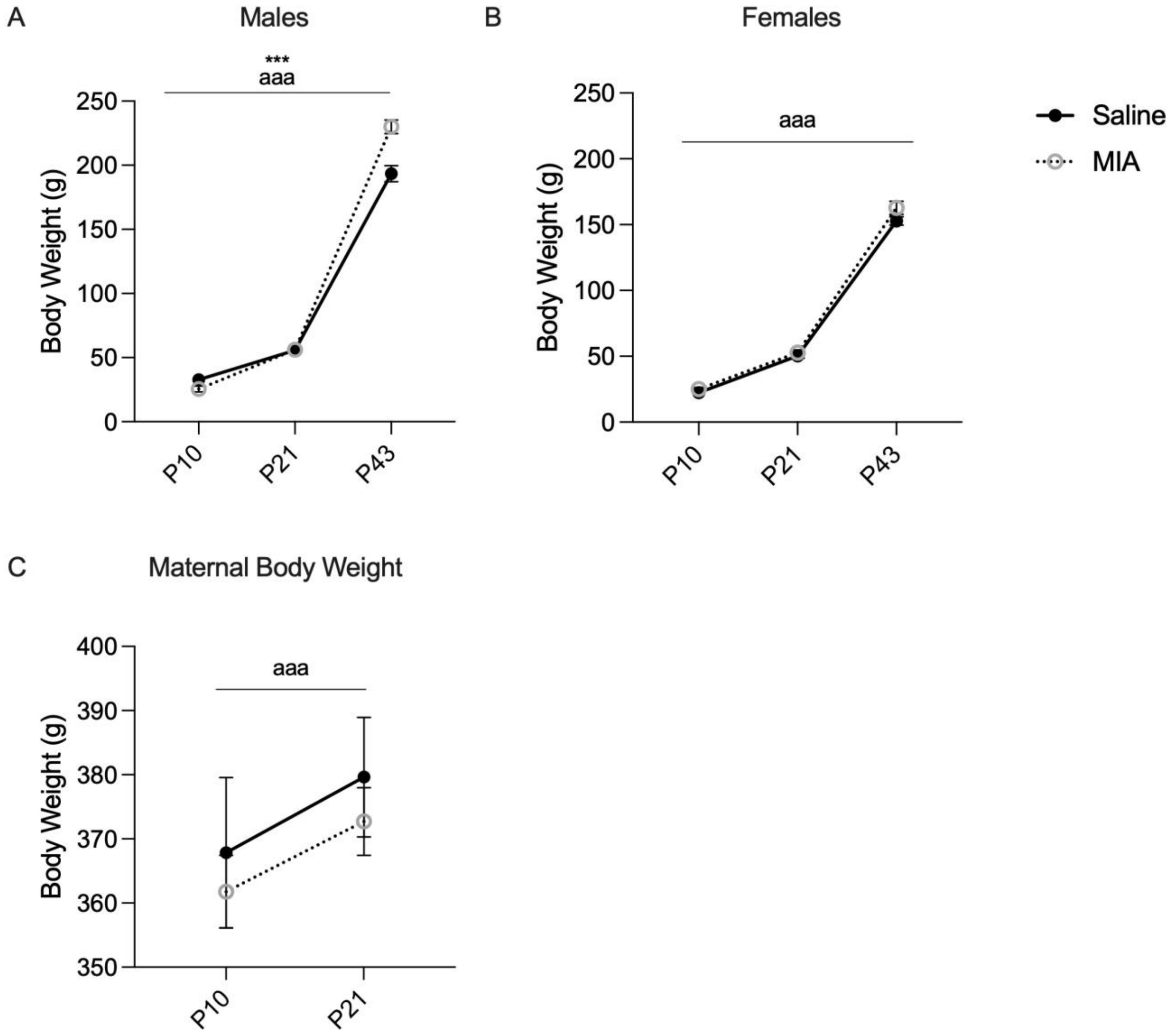


**Supplementary Figure 7. Offspring and maternal body weights across early development following maternal immune activation (MIA) during the mid-lactational period.** A) Male body weights demonstrated a significant effect of time ($F(2, 26)= 1078.59$, $p<0.001$) and MIA ($F(1,13)= 7618.25$, $p<0.001$) while B) female offspring and C) dam body weights only differed significantly across time (females: $F(2, 26)=1491.4$, $p<0.001$; maternal body weight: $F(1, 13)=21.64$, $p<0.001$). P10 offspring body weights were obtained by taking the average weight from the weight of all pups of the same sex for each litter (Saline: n = 8; MIA n = 7). P21 and P43 body weights were obtained from one male and one female per litter (Saline: n = 8; MIA n = 7). Data are expressed as mean ± SEM. ***$p < 0.001$, main effect of MIA; $^{aaa}p < 0.001$, main effect of time.

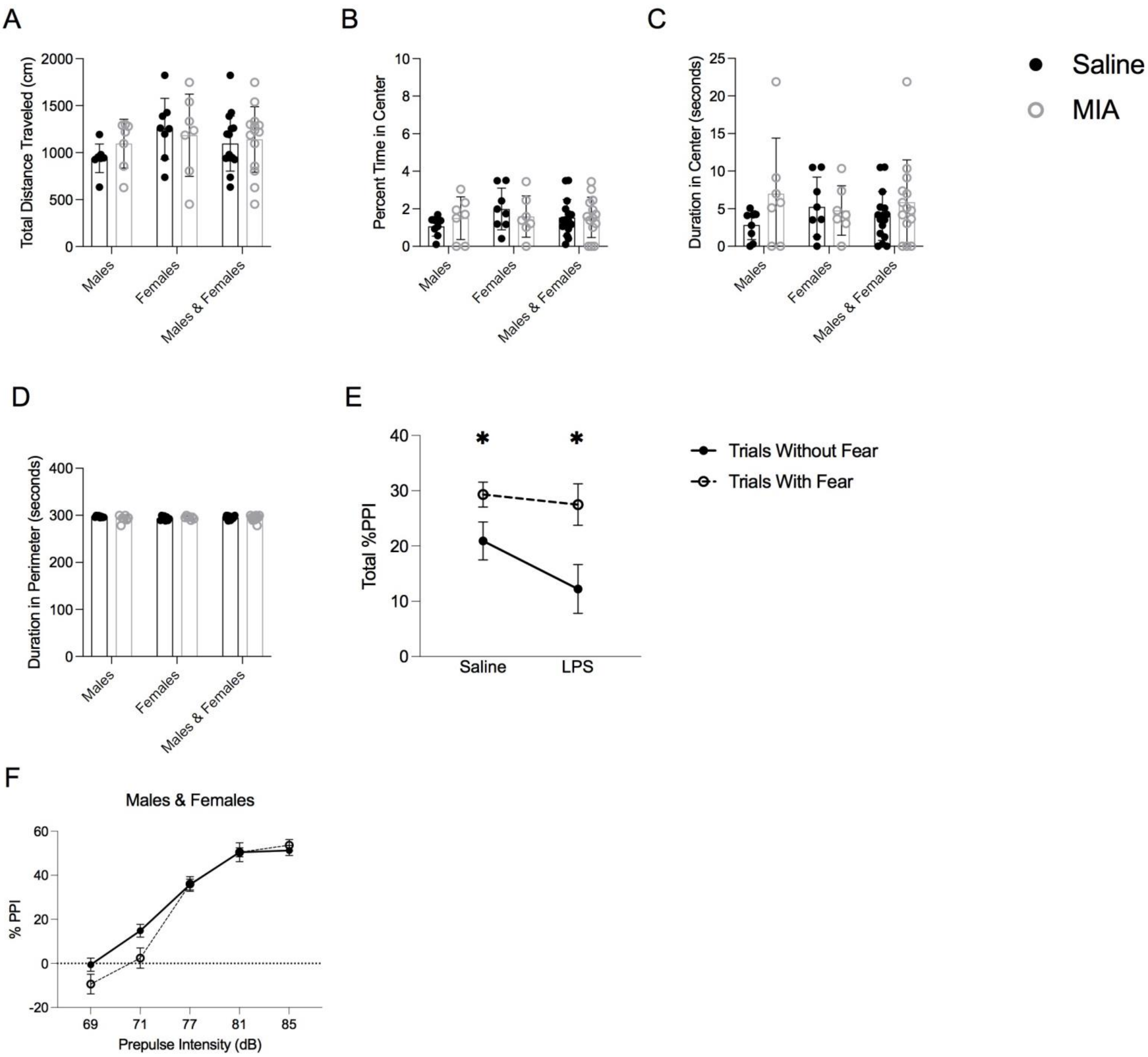


**Supplementary Figure 8. Adolescent offspring behavior following maternal immune activation (MIA) during the mid-lactational period.** (A) Distance traveled (cm), (B) percent time spent in the center, and duration of time (seconds) spent in the (C) center and (D) perimeter of an open field arena. (E) Conditioned fear significantly increased %PPI regardless of MIA or sex ($F(4, 52)= 8.19$, $p=0.008$, $n_p^2=0.226$). (F) %PPI for trials primed with conditioned fear collapsed across males and females for display purposes. Saline: n = 8; MIA n = 7. Data are expressed as mean ± SEM. Data are expressed as mean ± SEM, *p<0.05.
