## Supplementary Table 2 for "Got Milk? Maternal immune activation during the mid-lactational period affects nutritional milk quality and adolescent offspring sensory processing in male and female rats"

| **Supplementary Table 2.** Differentially expressed genes in P10 milk samples within the targeted pathways. | | | | |
|---|---|---|---|---|
| **Pathway** | **Gene/s** | **p-value** | **padj** | **FC** |
| Milk triglycerides, creamatocrit, and fatty acids | *Fads1* | 0.0405 | 0.481 | 0.365 |
| | *Elovl1* | 0.0270 | 0.420 | 0.399 |
| | *Abcg2* | 0.0211 | 0.378 | 0.450 |
| | *Igf1* | 0.0011 | 0.099 | 1.379 |
| Nutrient transport | *Slc5A1* | 0.0337 | 0.456 | 0.492 |
| | *Csn1S2A* | 0.0221 | 0.385 | 0.559 |
| | *Abcg2* | 0.0211 | 0.378 | 0.450 |
| | *Slc5A3* | 0.0001 | 0.007 | 1.329 |
| | *Tf* | 0.0001 | 0.001 | 1.669 |
| | *Csn1S2B* | 0.0001 | 0.001 | 1.733 |
| Oxytocin | *Jun* | 0.0015 | 0.112 | 1.007 |
| | *Fos* | 0.0038 | 0.170 | 1.956 |
| Aldosterone/MR | *Ace* | 0.0196 | 0.517 | 1.082 |
| | *Nr3c2* | 0.0001 | 0.013 | -0.828 |
| Epigenetics | *Kdm1a* | 0.0382 | 0.471 | -0.225 |
| Prolactin | *Jak2* | 0.0033 | 0.161 | 0.474 |
| | *Src* | 0.0263 | 0.418 | -0.604 |
| Parvalbumin | *Foxp2* | 0.0385 | 0.472 | 0.687 |
| | *Rnd3* | 0.0003 | 0.044 | 1.684 |
| | *Tenm3* | 0.0001 | 0.010 | -2.393 |
| | *Meis3* | 0.0001 | 0.006 | -1.985 |
| Glutamate/GABA | *Cpt1c* | 0.0116 | 0.278 | 0.861 |
| | *Slc7a11* | 0.0222 | 0.385 | 0.785 |
| | *Gabbr2* | 0.0241 | 0.403 | 1.358 |
| Inflammation | *Traf2* | 0.0172 | 0.341 | -0.322 |
| | *Myd88* | 0.0172 | 0.341 | 0.583 |
| | *Cfb* | 0.0242 | 0.403 | 0.686 |
| | *Tlr3* | 0.0384 | 0.471 | 0.644 |
| | *Tnfrsf1A* | 0.0476 | 0.495 | 0.359 |
| GR Binding | *Fam19a5* | 0.0001 | 0.006 | 1.453 |
| | *Nr3c2* | 0.0001 | 0.012 | -0.828 |
| | *Azin1* | 0.0003 | 0.054 | 0.407 |
| | *Ttyh1* | 0.0004 | 0.056 | 1.759 |
| | *Cndp1* | 0.0004 | 0.061 | 0.992 |
| | *Hpn* | 0.0006 | 0.069 | 0.703 |
| | *Lao1* | 0.0006 | 0.069 | 0.817 |
| | *St3gal4* | 0.0007 | 0.082 | -1.141 |

| Pathway | Gene/s | p-value | padj | FC |
|---|---|---|---|---|
| | *Mfge8* | 0.0008 | 0.088 | 0.648 |
| | *Ksr1* | 0.0010 | 0.098 | 1.315 |
| | *Pim3* | 0.0016 | 0.116 | -0.815 |
| | *Uspl1* | 0.0017 | 0.120 | -0.494 |
| | *Tmem140* | 0.0018 | 0.122 | 1.365 |
| | *Hspa1b* | 0.0023 | 0.142 | 0.799 |
| | *Ifi30* | 0.0026 | 0.153 | 0.402 |
| | *Wdr5* | 0.0028 | 0.156 | -0.346 |
| | *Utrn* | 0.0032 | 0.161 | -0.447 |
| | *Fos* | 0.0038 | 0.170 | 1.956 |
| | *Tsc22d1* | 0.0042 | 0.178 | 0.822 |
| | *Bambi* | 0.0045 | 0.186 | 1.650 |
| | *Bcor* | 0.0045 | 0.186 | 0.544 |
| | *Muc15* | 0.0050 | 0.194 | 0.652 |
| | *Urb1* | 0.0054 | 0.201 | -0.521 |
| | *Krt80* | 0.0054 | NA | 2.231 |
| | *Ncald* | 0.0055 | 0.201 | 0.864 |
| | *Egln3* | 0.0055 | 0.201 | -1.030 |
| | *Cst3* | 0.0061 | 0.211 | 0.791 |
| | *Atp6ap2* | 0.0064 | 0.215 | 0.497 |
| | *Hopx* | 0.0065 | 0.215 | 0.650 |
| | *Iqgap1* | 0.0068 | 0.217 | 0.493 |
| | *Cblb* | 0.0068 | 0.217 | -0.709 |
| | *Nampt* | 0.0069 | 0.219 | 0.387 |
| | *Fam180a* | 0.0071 | 0.222 | 0.955 |
| | *Parvg* | 0.0073 | NA | 1.857 |
| | *Arrdc2* | 0.0073 | 0.226 | 0.941 |
| | *Xpo6* | 0.0079 | 0.234 | -0.420 |
| | *Plce1* | 0.0079 | 0.234 | -0.447 |
| | *Ptpn9* | 0.0079 | 0.234 | -0.355 |
| | *Dagla* | 0.0079 | 0.234 | 1.305 |
| | *Chrdl2* | 0.0086 | 0.239 | -0.955 |
| | *Ogfrl1* | 0.0098 | 0.256 | 0.513 |
| | *Acsl3* | 0.0100 | 0.259 | 0.974 |
| | *Cadps* | 0.0102 | NA | -4.321 |
| | *Slc25a25* | 0.0106 | 0.267 | -0.678 |
| | *Gch1* | 0.0110 | 0.270 | 1.639 |
| | *Arhgef7* | 0.0123 | 0.288 | -0.299 |
| | *Nfkbiz* | 0.0134 | 0.306 | 1.508 |
| | *Irf2* | 0.0136 | 0.306 | 0.578 |

| Pathway | Gene/s | p-value | padj | FC |
|---|---|---|---|---|
| | *Sipa1l1* | 0.0136 | 0.306 | -0.482 |
| | *Coro1c* | 0.0143 | 0.314 | -0.444 |
| | *Rtn1* | 0.0146 | 0.318 | -1.000 |
| | *Car6* | 0.0147 | 0.319 | 0.628 |
| | *Capn2* | 0.0165 | 0.336 | -0.336 |
| | *Col5a1* | 0.0166 | 0.336 | 1.090 |
| | *Tmcc3* | 0.0166 | 0.336 | 0.659 |
| | *Gfod1* | 0.0168 | 0.336 | -0.687 |
| | *Baz2b* | 0.0171 | 0.341 | 0.483 |
| | *Nxt1* | 0.0178 | 0.350 | -0.386 |
| | *Tob2* | 0.0185 | 0.354 | -0.811 |
| | *Sdc4* | 0.0208 | 0.378 | 1.735 |
| | *Calml3* | 0.0214 | NA | 1.570 |
| | *Clcn7* | 0.0216 | 0.382 | 0.462 |
| | *Sulf2* | 0.0220 | 0.385 | 0.475 |
| | *Slc7a11* | 0.0222 | 0.385 | 0.785 |
| | *Mknk1* | 0.0227 | 0.390 | 0.399 |
| | *Gabbr2* | 0.0241 | 0.403 | 1.358 |
| | *Chst1* | 0.0245 | 0.406 | -0.808 |
| | *Sult5a1* | 0.0247 | 0.407 | 0.272 |
| | *Hsf2* | 0.0251 | 0.408 | -0.494 |
| | *Agpat4* | 0.0263 | NA | -3.138 |
| | *Abl2* | 0.0273 | 0.423 | -0.543 |
| | *Ankrd9* | 0.0280 | 0.426 | -0.592 |
| | *Igfbp7* | 0.0296 | 0.435 | 0.891 |
| | *Per1* | 0.0299 | 0.437 | -0.530 |
| | *Usp12* | 0.0303 | 0.439 | -0.280 |
| | *Chka* | 0.0312 | 0.445 | -0.426 |
| | *Sh2d4a* | 0.0315 | 0.447 | 0.509 |
| | *Ncam1* | 0.0317 | NA | 1.237 |
| | *Dnajc6* | 0.033 | 0.454 | -0.275 |
| | *Cerk* | 0.0335 | 0.456 | 0.507 |
| | *Dusp6* | 0.0340 | 0.457 | 1.218 |
| | *Epha2* | 0.0345 | 0.460 | 0.975 |
| | *Ap1m1* | 0.0349 | 0.461 | -0.320 |
| | *Tm2d1* | 0.0355 | 0.462 | 0.414 |
| | *Mgst2* | 0.0383 | 0.471 | 0.936 |
| | *Agtrap* | 0.0387 | 0.473 | 0.402 |
| | *Lyplal1* | 0.0392 | 0.475 | 0.344 |
| | *Lrrk1* | 0.0400 | NA | 2.008 |

| Pathway | Gene/s | p-value | padj | FC |
|---|---|---|---|---|
| | *Tmtc2* | 0.0412 | 0.485 | 0.467 |
| | *Cdkn2b* | 0.0413 | 0.485 | 0.817 |
| | *Thada* | 0.0418 | 0.485 | -0.330 |
| | *Rbl2* | 0.0425 | 0.486 | 0.338 |
| | *Bst1* | 0.0427 | NA | 2.197 |
| | *Nedd9* | 0.0438 | 0.488 | 0.879 |
| | *Tpk1* | 0.0445 | 0.490 | 0.295 |
| | *Hsd17b7* | 0.0452 | 0.492 | 0.638 |
| | *Sirpa* | 0.0461 | 0.494 | 0.405 |
| | *Ahdc1* | 0.0472 | 0.495 | -0.533 |
| | *Sat1* | 0.0472 | 0.495 | 0.346 |
| | *Klf3* | 0.0475 | 0.495 | 0.398 |
| | *Socs5* | 0.0475 | 0.495 | -0.291 |
| | *Tnfrsf1a* | 0.0476 | 0.495 | 0.359 |
| | *Pmepa1* | 0.0483 | 0.500 | -0.939 |
| | *Ucp3* | 0.0488 | 0.501 | -0.487 |
| | *Frmd4a* | 0.0496 | 0.502 | -0.820 |
| | *Cd81* | 0.0497 | 0.502 | 0.369 |
| | *Atxn10* | 0.0499 | 0.503 | -0.251 |
