## Supplementary Table 3 for "Got Milk? Maternal immune activation during the mid-lactational period affects nutritional milk quality and adolescent offspring sensory processing in male and female rats"

| **Supplementary Table 3.** Differentially expressed genes in P11 milk samples within the targeted pathways. | | | | |
|---|---|---|---|---|
| **Pathway** | **Gene/s** | **p-value** | **padj** | **FC** |
| Milk triglycerides, creamatocrit, and fatty acids | *Fto* | 0.0156 | 0.497 | 0.378 |
| Nutrient transport | *Tf* | 0.0054 | 0.344 | 0.904 |
| Oxytocin | *Cacnb1* | 0.0169 | 0.509 | 4.128 |
| | *Ryr2* | 0.0254 | 0.548 | 4.838 |
| | *Itpr1* | 0.0429 | 0.619 | 0.623 |
| | *Jun* | 0.0466 | 0.628 | -0.336 |
| LPS Binding | *Irf5* | 0.0233 | 0.542 | -1.267 |
| Glucocorticoid Signaling | *Dnajb1* | 0.0462 | 0.628 | -0.368 |
| Epigenetics | *Kdm6b* | 0.0043 | 0.318 | -0.769 |
| | *Hdac3* | 0.0064 | 0.376 | 0.269 |
| | *Smarcal1* | 0.0131 | 0.472 | 0.534 |
| | *Chd2* | 0.0189 | 0.517 | -0.410 |
| | *Slc23a2* | 0.0237 | 0.543 | -0.534 |
| | *Kdm3a* | 0.0251 | 0.547 | -0.306 |
| Prolactin | *Prlr* | 0.0373 | 0.601 | 0.869 |
| Parvalbumin | *Kcng4* | 0.0154 | 0.492 | -0.607 |
| | *Wnt5A* | 0.0224 | 0.538 | 1.509 |
| | *Wnt5B* | 0.0382 | 0.608 | 1.027 |
| Glutamate/GABA | *Gria3* | 0.0120 | 0.467 | -1.858 |
| | *Pias1* | 0.0283 | 0.568 | -0.454 |
| | *Slc1a3* | 0.0309 | 0.581 | 4.576 |
| | *Gpsm1* | 0.0384 | 0.608 | -0.722 |
| Inflammation | *Stat2* | 0.0009 | 0.156 | -0.569 |
| | *Cfb* | 0.0012 | 0.181 | 0.910 |
| | *Tlr3* | 0.0053 | 0.341 | 0.608 |
| | *Tnfrsf1B* | 0.0077 | 0.402 | -1.92 |
| | *C3* | 0.0156 | 0.497 | 0.784 |
| | *C1Qa* | 0.0164 | 0.501 | 1.352 |
| | *Myd88* | 0.0236 | 0.542 | -0.343 |
| | *Jak3* | 0.0269 | 0.556 | 0.600 |
| | *Il6R* | 0.0334 | 0.589 | -0.522 |
| | *Il1B* | 0.0431 | 0.620 | -1.693 |
| | *C2* | 0.0443 | 0.624 | 0.484 |
| | *Tnfrsf1A* | 0.0446 | 0.625 | -0.343 |
| GR Binding | *Lao1* | 0.0002 | 0.066 | 1.402 |
| | *Zfp36* | 0.0005 | 0.120 | -0.806 |
| | *Shc4* | 0.0005 | 0.120 | -1.340 |

| Pathway | Gene/s | p-value | padj | FC |
|---|---|---|---|---|
| | *Mafk* | 0.0005 | 0.123 | -0.654 |
| | *Cst3* | 0.0007 | 0.148 | 0.507 |
| | *Cryab* | 0.0010 | 0.164 | 0.854 |
| | *Fam84a* | 0.0012 | 0.185 | 0.800 |
| | *Il17rd* | 0.0015 | 0.206 | 1.459 |
| | *Tsc22d3* | 0.0015 | 0.206 | -0.579 |
| | *Enpp4* | 0.0017 | 0.214 | -2.302 |
| | *Eif2ak2* | 0.0019 | 0.231 | -0.819 |
| | *Gadd45g* | 0.0028 | 0.268 | -0.464 |
| | *Etnk2* | 0.0028 | 0.270 | -3.064 |
| | *Gpbp1* | 0.0033 | 0.291 | -0.336 |
| | *Decr1* | 0.0035 | 0.292 | 0.622 |
| | *Alcam* | 0.0035 | 0.293 | -0.511 |
| | *Aven* | 0.0037 | 0.300 | 0.607 |
| | *Rbm46* | 0.0042 | 0.315 | 0.902 |
| | *Kdm6b* | 0.0043 | 0.318 | -0.769 |
| | *Nfkbiz* | 0.0047 | 0.329 | -0.732 |
| | *Cachd1* | 0.0057 | 0.356 | 1.811 |
| | *Tle4* | 0.0065 | 0.376 | -0.509 |
| | *Slc25a4* | 0.0067 | 0.384 | 1.151 |
| | *Glud1* | 0.0071 | 0.397 | 0.342 |
| | *Cpe* | 0.0076 | 0.402 | -2.449 |
| | *Tmem140* | 0.0076 | 0.402 | -0.766 |
| | *Clk3* | 0.0076 | 0.402 | -0.367 |
| | *Ifih1* | 0.0080 | 0.407 | -1.328 |
| | *B4galnt1* | 0.0083 | 0.410 | 4.148 |
| | *Nfil3* | 0.0086 | 0.410 | -0.356 |
| | *Nfkbia* | 0.0093 | 0.421 | -0.656 |
| | *Hspa1a* | 0.0099 | 0.438 | -0.688 |
| | *Nubpl* | 0.0100 | 0.440 | 0.549 |
| | *Hopx* | 0.0101 | 0.440 | -0.679 |
| | *Myom1* | 0.0101 | 0.440 | -0.699 |
| | *Ypel5* | 0.0103 | 0.441 | -0.343 |
| | *Raver2* | 0.0112 | 0.456 | 4.329 |
| | *Galnt10* | 0.0122 | 0.467 | 0.591 |
| | *Arrdc2* | 0.0123 | 0.467 | -0.498 |
| | *Usp25* | 0.0134 | 0.474 | -0.505 |
| | *Prlhr* | 0.0136 | 0.475 | -2.409 |
| | *Ppara* | 0.0145 | 0.488 | 1.010 |
| | *Ralgds* | 0.0145 | 0.488 | -0.498 |

| Pathway | Gene/s | p-value | padj | FC |
|---|---|---|---|---|
| | *Sulf2* | 0.0146 | 0.488 | 0.575 |
| | *Notch2* | 0.0147 | 0.488 | -0.500 |
| | *Fto* | 0.0156 | 0.497 | 0.378 |
| | *Spsb1* | 0.0159 | 0.499 | 1.329 |
| | *Fam126a* | 0.0162 | 0.501 | -0.761 |
| | *Baz2b* | 0.0179 | 0.516 | -0.385 |
| | *Fxyd3* | 0.0181 | 0.516 | 0.641 |
| | *Mapk10* | 0.0181 | 0.516 | -1.445 |
| | *Hsf2* | 0.0187 | 0.517 | -0.456 |
| | *Ppp1r1b* | 0.0188 | 0.517 | 0.828 |
| | *Luzp1* | 0.0199 | 0.517 | -0.368 |
| | *Arhgap24* | 0.0201 | 0.517 | -1.631 |
| | *Pcgf5* | 0.0204 | 0.520 | -0.459 |
| | *Dolk* | 0.0211 | 0.528 | 0.499 |
| | *Adamtsl4* | 0.0214 | 0.530 | -0.779 |
| | *Amz2* | 0.0220 | 0.533 | 0.396 |
| | *Atm* | 0.0231 | 0.542 | 0.419 |
| | *Nuak2* | 0.0244 | 0.544 | -1.053 |
| | *Mgst2* | 0.0246 | 0.544 | 0.747 |
| | *Agpat3* | 0.0256 | 0.551 | 0.405 |
| | *Gucy2g* | 0.0257 | 0.552 | 0.912 |
| | *Impa1* | 0.0266 | 0.556 | 0.308 |
| | *Hspa1b* | 0.0267 | 0.556 | -0.550 |
| | *Gna13* | 0.0272 | 0.558 | -0.442 |
| | *Tnfaip2* | 0.0277 | 0.565 | -1.292 |
| | *Chka* | 0.0282 | 0.568 | -0.252 |
| | *Socs5* | 0.0284 | 0.568 | 0.460 |
| | *Trmu* | 0.0292 | 0.573 | 0.410 |
| | *Lrsam1* | 0.0301 | 0.575 | 0.453 |
| | *Ptpru* | 0.0303 | 0.577 | 4.799 |
| | *Slc1a3* | 0.0309 | 0.581 | 4.576 |
| | *Nrxn1* | 0.0323 | 0.585 | 4.205 |
| | *Depdc7* | 0.0327 | 0.587 | -0.447 |
| | *Mpzl3* | 0.0329 | 0.587 | -0.427 |
| | *Parva* | 0.0331 | 0.587 | 0.393 |
| | *Tmtc2* | 0.0335 | 0.590 | 0.799 |
| | *Mbp* | 0.0337 | 0.592 | -0.484 |
| | *Lrch1* | 0.0340 | 0.592 | -0.301 |
| | *Ell* | 0.0342 | 0.592 | -0.463 |
| | *Stk17b* | 0.0344 | 0.592 | -0.494 |

| Pathway | Gene/s | p-value | padj | FC |
|---|---|---|---|---|
| | *Tbx4* | 0.0344 | 0.592 | -1.967 |
| | *Cdyl* | 0.0345 | 0.592 | -0.434 |
| | *Slc15a2* | 0.0348 | 0.596 | 1.116 |
| | *Etnk1* | 0.0351 | 0.596 | -0.452 |
| | *Irak2* | 0.0358 | 0.596 | -0.678 |
| | *Fgf2* | 0.0359 | 0.598 | 3.759 |
| | *Slc22a23* | 0.0370 | 0.601 | 0.355 |
| | *As3mt* | 0.0371 | 0.601 | 0.452 |
| | *Mdga1* | 0.0372 | 0.601 | -1.199 |
| | *Tgfbr3* | 0.0381 | 0.608 | 0.627 |
| | *Wnt5b* | 0.0382 | 0.608 | 1.027 |
| | *Gpsm1* | 0.0384 | 0.608 | -0.722 |
| | *Nrxn3* | 0.0394 | 0.611 | 3.231 |
| | *Ptpn1* | 0.0394 | 0.611 | -0.280 |
| | *Lgr4* | 0.0395 | 0.611 | 0.381 |
| | *Cdh23* | 0.0399 | 0.612 | -1.746 |
| | *Ndfip2* | 0.0408 | 0.614 | -0.314 |
| | *Lmod1* | 0.0410 | 0.614 | -0.709 |
| | *Atg10* | 0.0412 | 0.614 | 0.798 |
| | *Rab31* | 0.0413 | 0.614 | -0.892 |
| | *Cd81* | 0.0417 | 0.614 | 0.257 |
| | *Hddc3* | 0.0418 | 0.615 | 0.645 |
| | *Lrrfip1* | 0.0423 | 0.617 | -0.721 |
| | *Itpr1* | 0.0429 | 0.619 | 0.623 |
| | *Tmem17* | 0.0433 | 0.620 | 0.677 |
| | *Oaf* | 0.0441 | 0.624 | -1.213 |
| | *Tnfrsf1a* | 0.0446 | 0.625 | -0.343 |
| | *Amotl2* | 0.0467 | 0.628 | -0.387 |
| | *Zmynd8* | 0.0469 | 0.628 | 0.420 |
| | *Lhpp* | 0.0487 | 0.629 | 0.460 |
| | *Ncald* | 0.0492 | 0.631 | 0.573 |
